# *De novo* design of anti-variant COVID-19 Vaccine

**DOI:** 10.1101/2022.10.20.513049

**Authors:** Arpita Goswami, S. Madan Kumar, Samee Ullah, Milind M. Gore

**Affiliations:** Leo’s Research Services and Suppliers, 171, Basaveshwaranagara, Hebbal, Mysuru 570016, India; Department of Chemistry-BMC Biochemistry, University of Uppsala, Husargatan 3, 75237, Uppsala, Sweden; National Centre for Bioinformatics (NCB), Islamabad, 45320, Pakistan; 5/1B, Krutika Co-Op Housing Society, Tejas Nagar, Kothrud, Pune 411039, India

**Keywords:** De-novo protein design, SARS-CoV-2 variants, Vaccine design, Molecular dynamics simulations, AlphaFold, RosettaFold, Virology

## Abstract

Recent studies have shown the efficacy of hybrid SARS-COV-2 vaccines using wild-type nucleocapsid (N) and Spike (S) protein. We upgraded this strategy one step further using clinically proven spike protein by considering the delta and post-delta omicron variant of concern (VOC) mutations and nucleocapsid peptides conferring T-cell immunity. Nucleocapsid peptides are considered much better immunological replacement of nucleocapsid proteins. Hence, peptide linking strategy is applied which is more economic for cellular biosynthesis than whole proteins. An envelope peptide with potent T-cell immune response is also selected. All these peptides are clustered in this hybrid spike’s designed cytoplasmic region separated by non-immunogenic helical linkers. The resulting domain is more folded in the construct devoid of transmembrane domain after AlphaFold analysis. Alongside, we also propose the idea of introduction of any T-cell peptide like other Human Corona Viruses (HuCoV) in these linker regions whenever required. In addition to SARS-COV-2, the same approach can be applied for any emergency or even long-term unsolved outbreaks of Influenza, Dengue and West Nile Virus etc. In this era of novelty as presented by subunit and nucleic acid vaccines, multiepitope strategies like this can help to combat multiple diseases successfully in real time to give hope for better future.

## Introduction

SARS-COV-2-19 has brought the world to an unprecedented standstill for the last four years ^1^. The massacre and chaos created are prevailing still. This infection demonstrates a wide clinical spectrum, from asymptomatic, mild, moderate disease to even long term positivity and mortality ^2, 3^. Also, new variants are appearing regularly with mutations in anti-spike antibody binding sites causing new panic ^4, 5^. Although, mortality in infected patients is lower than earlier SARS, serious further complications are seen in reinfected and even in vaccinated populations. These include myocarditis, damage to CNS etc. that too in 30-40 years’ population ^6^. Reinfection with emerging newer variants is also a cause of concern ^7–9^.

In this scenario question arises on sustainability of vaccines based on only wild type spike ^10^. The spike protein which elicits neutralizing antibodies is essential for recovery ^11^. However, the problem is the short term of the direct mucosal immune response. Long-term memory in mucosal immune response is generated by T cell response ^12 13^. In children older than 2 years, infections by closely related corona viruses (Common cold) generate mild and asymptotic response against OC43, HKU1, 229E, NL63 (HuCoV) ^14 15^. Several immune response studies on these HuCoV infections have been carried out ^16 17 18^ so far. By the age of 2-5 years, 80% of children are immunized and have persistent neutralizing antibodies for life ^16^. The associated T cell response is also highly effective to generate memory response and lower severity in infected COVID patients ^19 20 21 22^. Several studies also suggest that T-cell immune response inducing nucleocapsid protein has to be included in potential corona vaccine strategy pipelines ^23 24 25 26^. This can be a promising approach to curb circulating Delta ^27^, Omicron variants ^28^ and other recombinants ^29^. Both Delta and Omicron (BA.1) are crucial as they mark the transition between pre-vaccination and post vaccination immune responses. Therefore, it is important to design a vaccine with delta-omicron spike protein and T cell epitopes. For this, we have designed a construct using chimeric SARS-COV-2 spike and SARS-COV-2 T cell epitope sequences. Structural proteins with almost conserved sequences for different corona virus families were targeted for potential T-cell enticing candidates. Functional proteins were not considered since some of them are known to hijack MHC systems in human ^30 31 32^. Several studies previously showed the homology of nucleocapsid protein sequences between SARS-COV-2 and HuCoV strains ^33 32 34 35^. Also, T-cell response of SARS-COV-2 unexposed individuals to SARS-COV-2 was previously reported for one peptide from envelope protein ^36^ indicating potential effect from common cold immunity. Therefore, we exclusively focused on nucleocapsid protein and envelope protein for T cell epitopes and compared with convalescent patients’ T cell response inducing sequences from literature. Also, more attention was given to CD4+ TCR epitopes than CD8+. Since the former sequences were found to be more stable than the other as per studies on patients ^37^. The designed sequence had 5 epitopes joined by linkers as cytoplasmic region. Among these, at least two epitopes were found folded after model building for the sequence without trans-membrane domain. As more well-formed structural epitopes can ensure more CD4+ T-cell activity ^38^ to cause persistent immunity. Lastly the structure was validated using the molecular dynamics simulation of 100ns time. Thus, we propose that this sequences and structures provided by our strategy could be quite useful and should be considered for vaccine design and development against SARS-COV-2 and can also be tweaked to any other future variants or viruses.

## Materials and Methods

The complete workflow of the current study is provided in figure 1.

**Figure 1.**
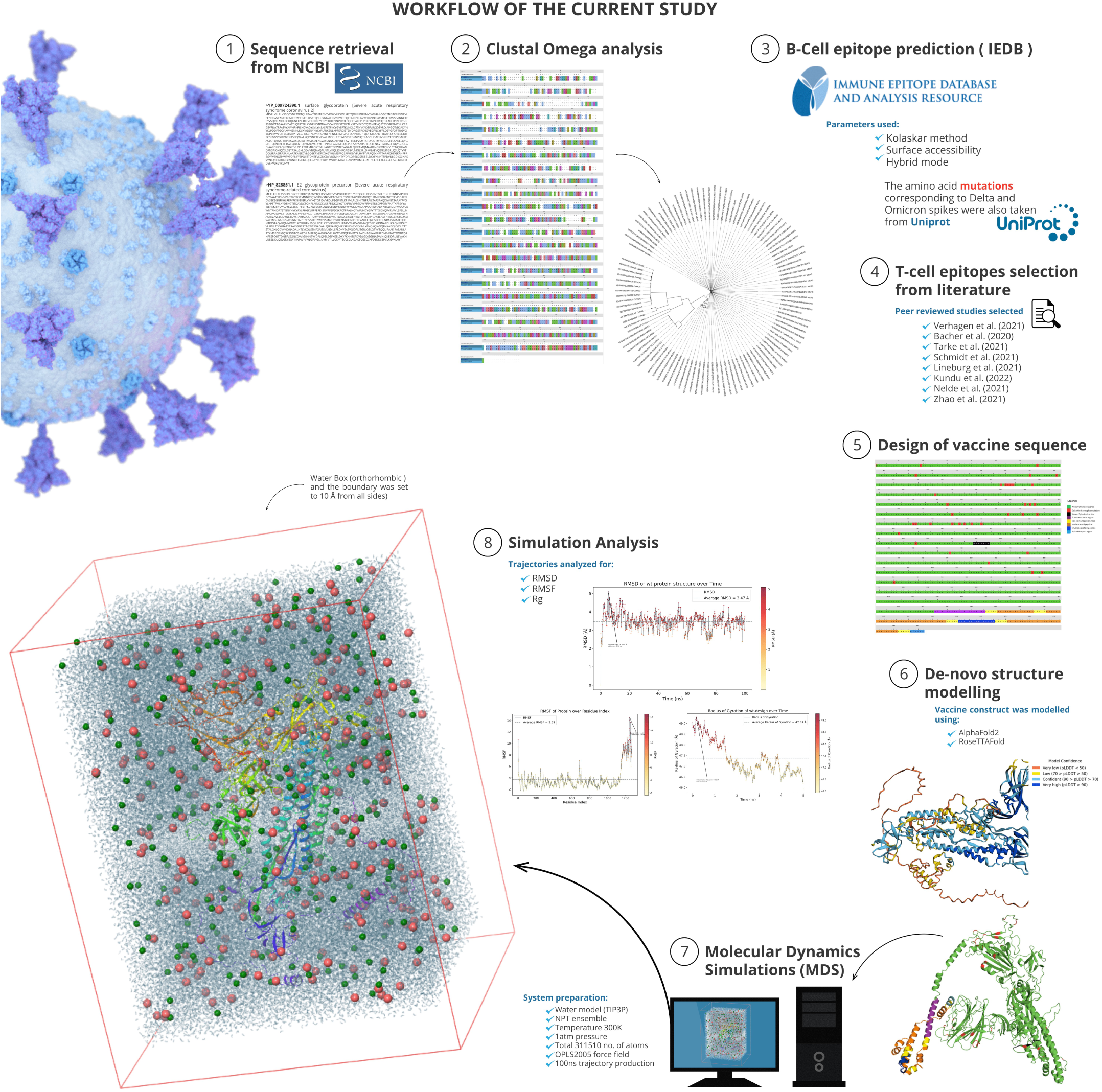
Workflow of the current study

## Sequences retrieval

The *SARS-1* and *SARS-CoV-2* spike sequences were retrieved from the UniProt database with accession no. P59594 and P0DTC2^39^. Sequences for *SARS-CoV-2* other structural proteins such as nucleocapsid (P0DTC9), Envelope protein (P0DTC4) and membrane protein (P0DTC5) were similarly acquired from Uniprot. In parallel, homologous structural protein sequences for HuCoV were also obtained along with *SARS-1*.

## Clustal Omega analysis

These sequences were imported into the Clustal Omega server of EBI for sequence alignment using the default protocol of ClustalW with characters count and before running the alignment by changing to maximum values for parameters such as guide tree iterations, HMM iterations and combined iterations (default value -1) ^40^.

## B-cell epitopes prediction

The *SARS-1* and *SARS-CoV-2* spike sequences were then imported into the (IEDB-Immune Epitope Database) server, and the B-cell epitopes predicted ^41^. The predictions parameters were set to Kolaskar method, surface accessibility and a hybrid mode. The predicted B-cell epitopes were color coded according to the method used in (Fig. 4). The individual protein domains were also mapped on the aligned sequences using information from both clustal alignment and UniProt. The amino acid mutations corresponding to Delta and Omicron spikes were also taken from Uniprot. All these mutations are substituted in native SARS-COV-2 spike sequence to design hybrid spike. The native furin site (PRRAR) is kept since it is a hotspot for immune response (from Uniprot).

## T cell epitopes selection from literature

The T cell epitopes from SARS-COV-2 recovered convalescent patients’ peer reviewed studies were selected from available extensive literatures ^17, 18, 20, 31, 32, 34, 40, 41^. Because the CD8+ epitopes cause nonpersistent immunity, in this study the literature was searched for CD4+ epitopes ^37^ and the potent epitopes with best possible folds of secondary structures as per PDBsum selected. Nucleocapsid and envelope peptides were preferred including one memory peptide from the former (with both T-cell and B-cell immunity). The similarities with homologous proteins from common cold coronavirus families as per literature studies were also included. The peptides having similar fold to human proteins structures including the closely situated in neighboring regions in individual open reading frames (ORFs) are excluded from this study to avoid allergenicity ^44^. Finally, to prevent misfolding in the design, the secondary structures of the selected peptides were predicted using the PDBsum and in addition the Phyre2 ^45, 46^.

## Sequence conservation analysis

Clustal omega sequence alignments for these proteins were generated using UniProt data on SARS-CoV-2, SARS, and common cold coronavirus. The alignment was refined by removing gapped sequences, non-human host harboring coronavirus homologs and partial protein fragments. The final alignment and associated sequence similarity-based neighbor joining tree (without distance corrections) were generated with maximum 5 iterations. Finally, the guide tree was imported and plotted in iTOL phylogenetic tree visualization server (interactive Tree of Life, ver. 6.5.8, EMBL) ^47^.

## Design of vaccine sequence

The hybrid sequence had mutated amino acids from both Delta and Omicron (BA.1) spikes. The core β strand was deleted from S2 region to prevent pre-to post fusion transition as shown before for coronavirus spike transitions (Fig. 6, 43). Potential Antibody dependent cellular cytotoxicity/Antibody dependent enhancement (ADCC/ADE) segment ^48–50^ mapped from Clustal O (Fig. 4) was further fine-tuned by structural alignment with *SARS* spike in PyMOL (Fig. 5). The minimum region stabilizing the same was deleted after verifying in AlphaFold-disorder package^51^. The N-epitopes interconnected by non-immunogenic EAAAK linkers were incorporated into the cytoplasmic domain ^52–55^. These epitopes were incorporated based to their native position in the nucleocapsid structure. The envelope peptide (15 residues) is placed between linkers among them to help in interaction (to cancel zero neighbors interaction effect from rigid EAAAK linker helices). The interaction of the last two N-epitopes with each other in PDB structure can be seen in the (Supplementary Fig. S6).

**Fig 1:**
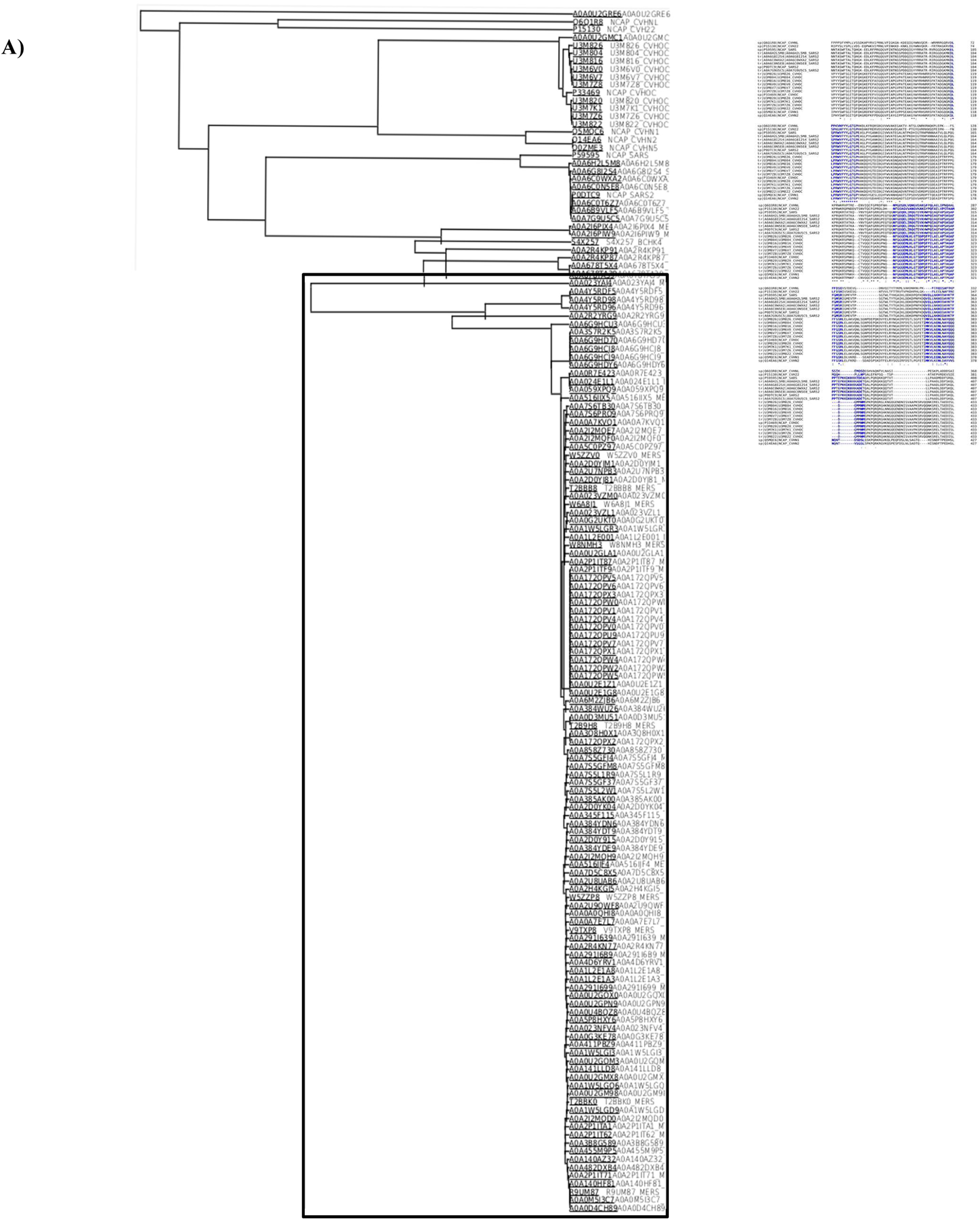
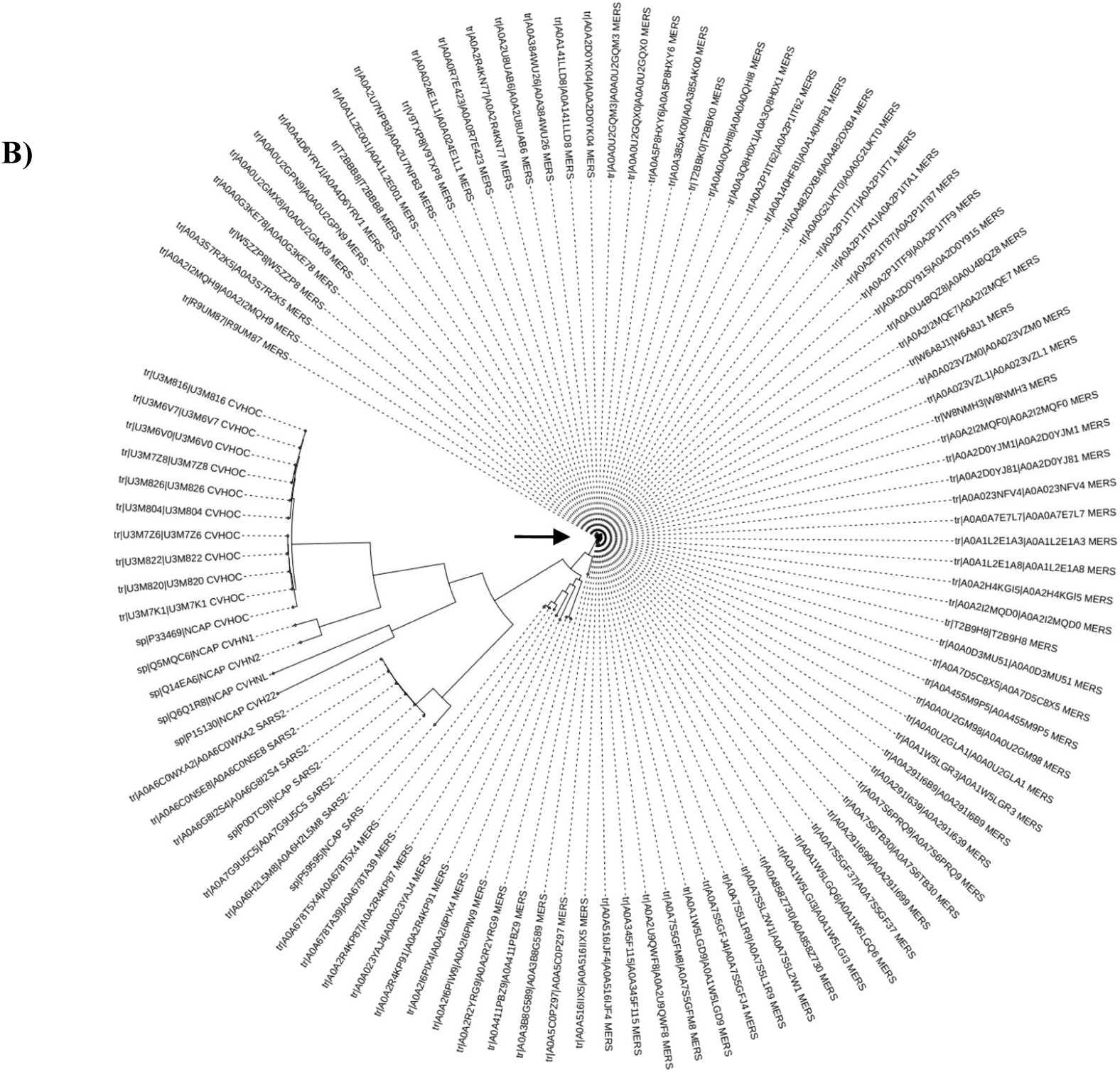
Clustal Omega alignment of Nucleocapsid (N) protein sequences. The sequence alignment among SARS, COVID and common cold coronaviruses shows high similarity in chosen epitopes (blue). Except last several residues in last peptide, all SARS2 N-epitopes are highly similar to HuCoV sequences.

**Fig 2:**
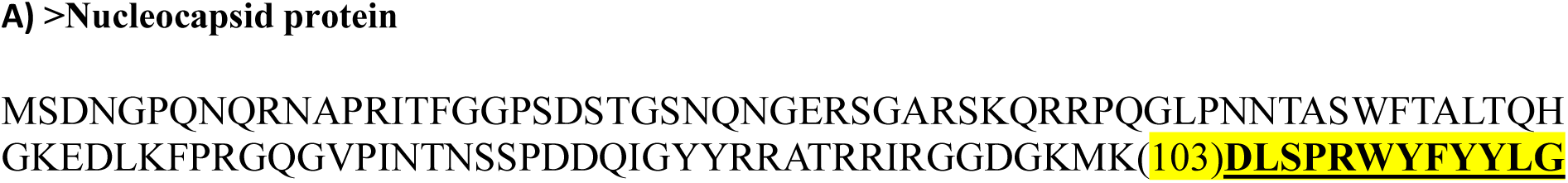
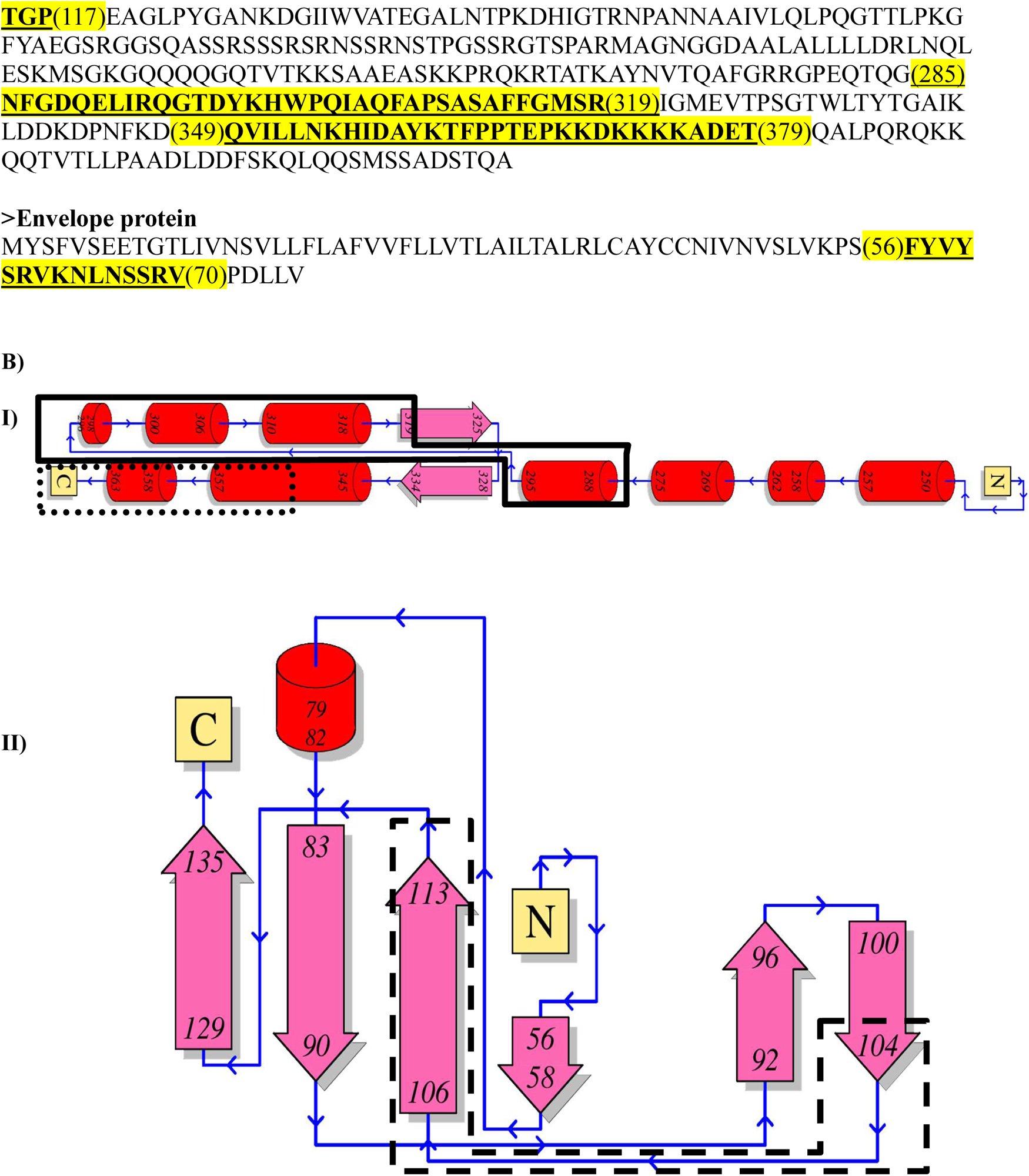
Sequence similarity-based guide tree of Coronavirus nucleocapsid sequences. **A)** Preliminary guide tree from Clustal Omega alignment of nucleocapsid sequences. Uniprot-retrieved Corona virus N-protein sequences’ alignment demonstrates distinct unique tree formation from later variants of MERS (box enclosed). Here all sequences as available from the database are used. **B)** Sequence similarity-based neighbouring-joining circular tree of coronavirus nucleocapsid proteins sequences. These are chosen from Uniprot after removing fragments, gapped sequences and non-human host infecting coronaviruses. For SARS, only one sequence was found to be without gaps. The circular tree is plotted in iTOL. Arrow marked are MERS nucleocapsid proteins showing very little changes in their sequences during as compared to others.

**Fig 3:**
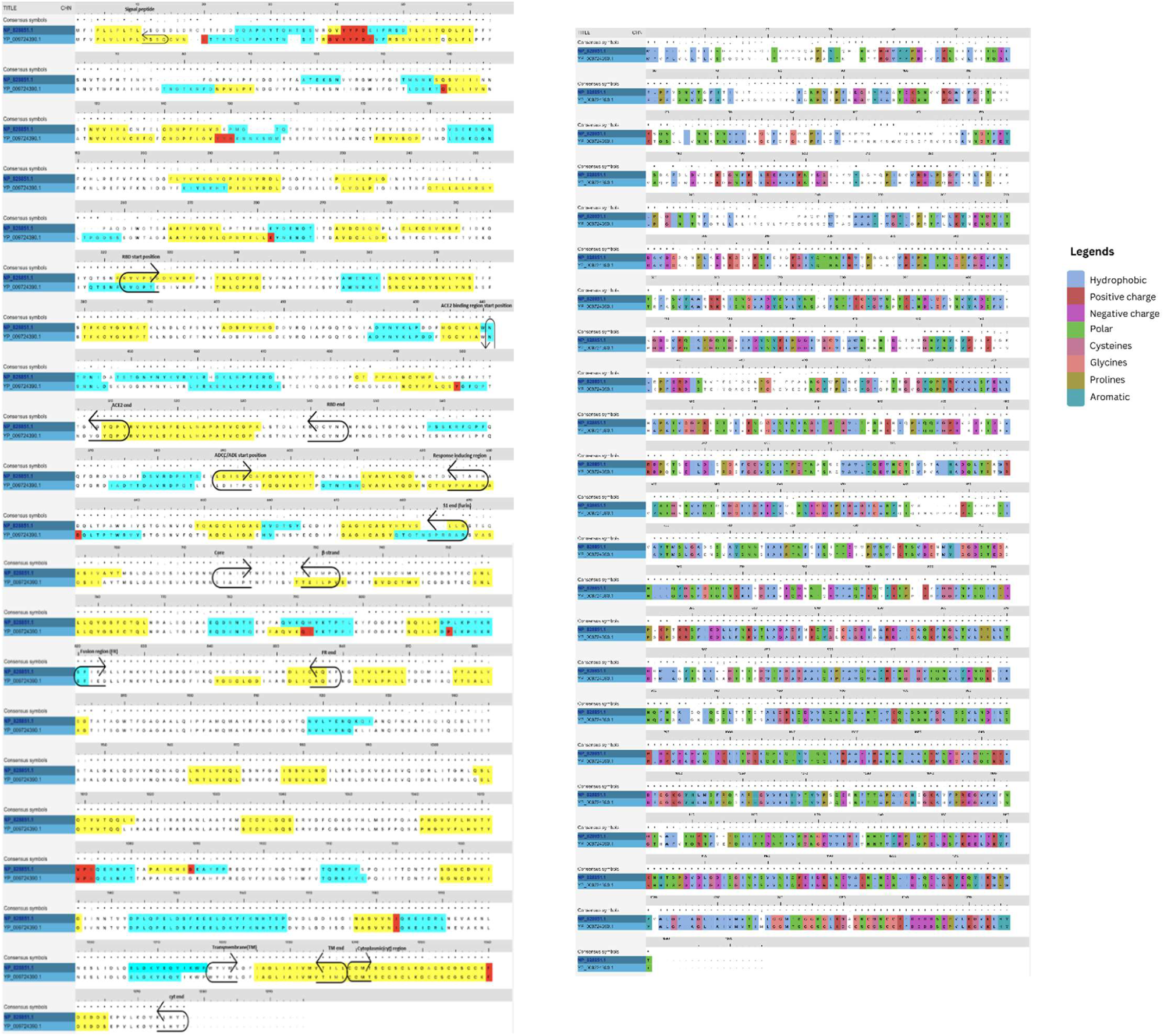
Wuhan COVID Nucleocapsid and Envelope protein epitopes. (A) The T-cell epitopes from N and E protein sequences (in FASTA format), homologous with common cold, chosen for the cytoplasmic domain design of the vaccine construct. These are highlighted yellow, bold-lettered and underlined. The numbers in parenthesis denote amino acid positions. (B) Secondary structures of the chosen nucleocapsid peptides in enclosures from PDBsum: I) **QVILLNKHIDAYKTFPPTEPKKDKKKKADET** (**6ZCO.pdb)** in dotted rectangle has mostly α-helical structure with adjoining loops; 15 residues missing at C terminus. **NFGDQELIRQGTDYKHWPQIAQFAPSASAFFGMSR (6ZCO.pdb)** in continuous lined enclosure has α-helical structures with adjoining loop. II) **DLSPRWYFYYLGTGP** (**7N0R.pdb)** in dashed enclosure has non-complementary β-strands along with adjoining long loop thereby strong possibility of forming unstructured epitope in construct design. The core sequence in this epitope is **SPRWYFYYL** which induces CD8 response (Supplemental Table 1). CD8+ epitopes are usually linear/unstructured.

**Fig 4:**
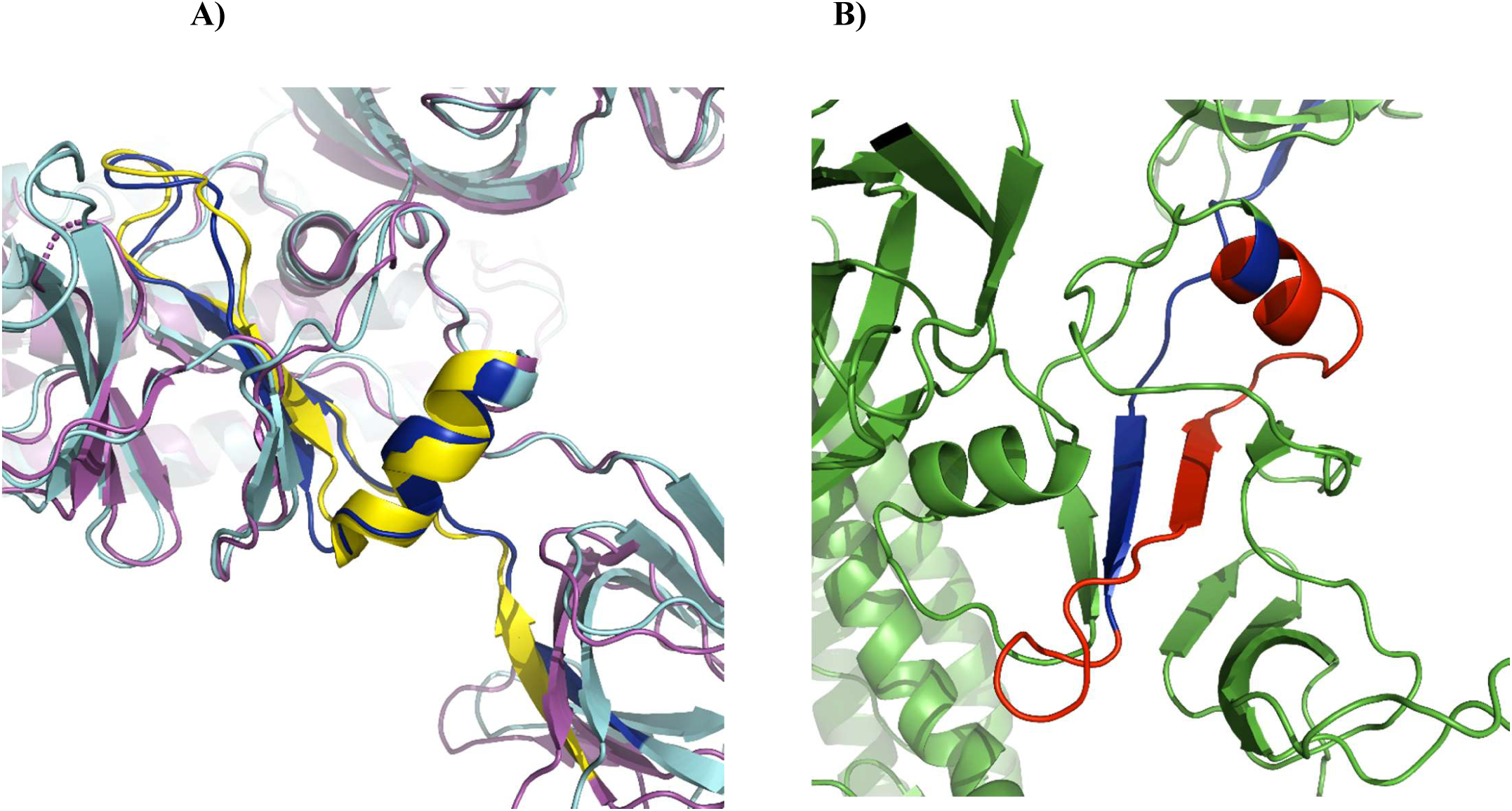
Mapping of B cell epitopes regions of Wuhan COVID spike (YP_009724390.1). Native SARS2 spike RBD (Receptor binding domain), ACE2 binding sequence, furin site, fusion region (FR) along with ADCC/ADE inducing regions are aligned with SARS spike (NP_828851.1) in Clustal Omega. The B-cell epitope analysis is from IEDB server. Yellow is B cell epitopes by Kolaskar method, Cyan blue is surface accessibility. Red is having both methods. These B cell epitope regions are present in delta and omicron spike also with ever-changing sequences evading immunity.

**Fig 5:**
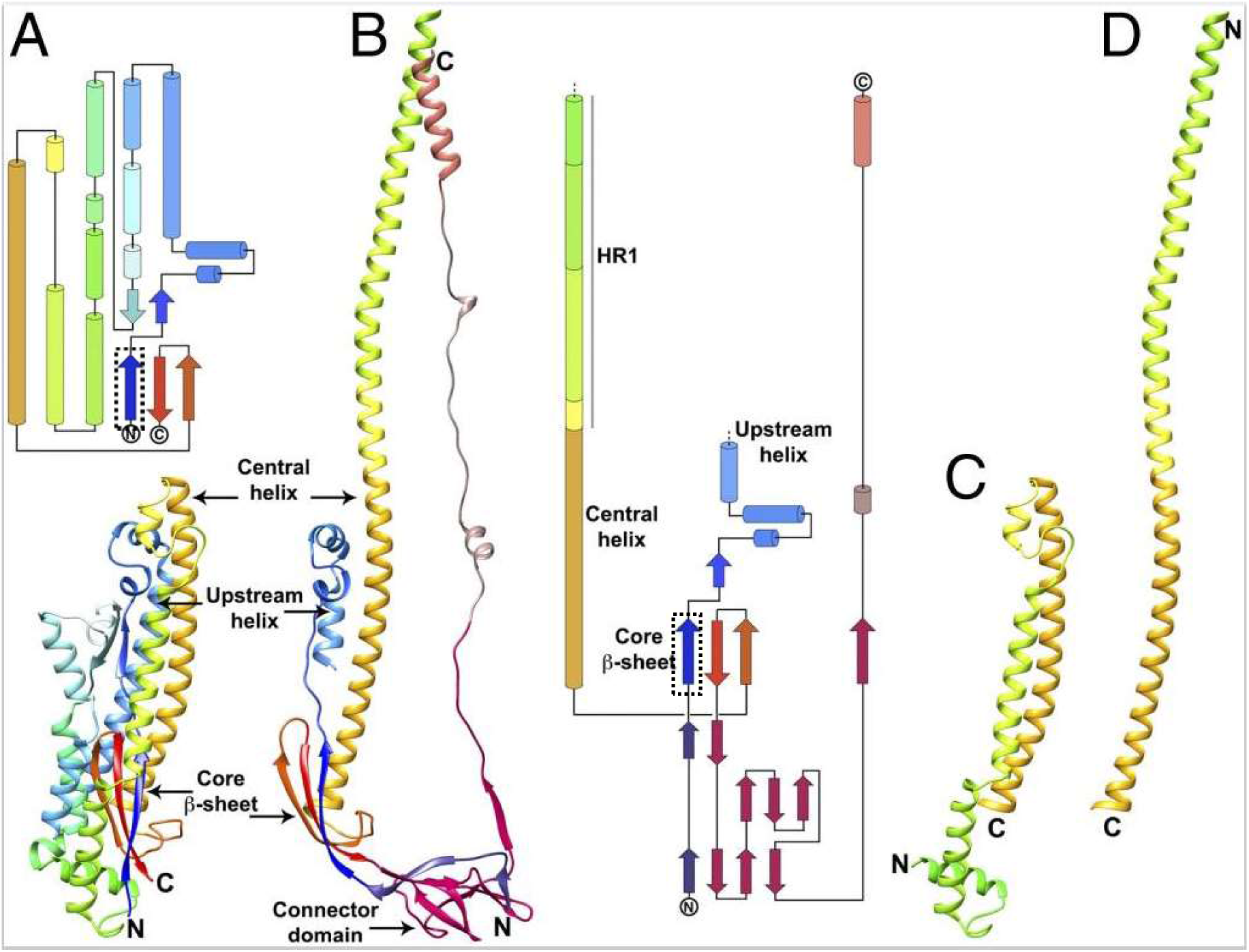
Mapping of ADCC and ADE region on COVID spike in PyMOL. **A)** Clustal O mapped region (585-625; blue) for COVID spike (cyan) is investigated by structural alignment on ADE inducing region (yellow) in SARS Spike (magenta). **B)** In vaccine construct designed, red segment (Δ600-624) was deleted as that will sufficiently weaken that region (rather a beta cage to encase any possible immunological proteins). Other non-ADE sequences are coloured as green.

**Fig 6:**
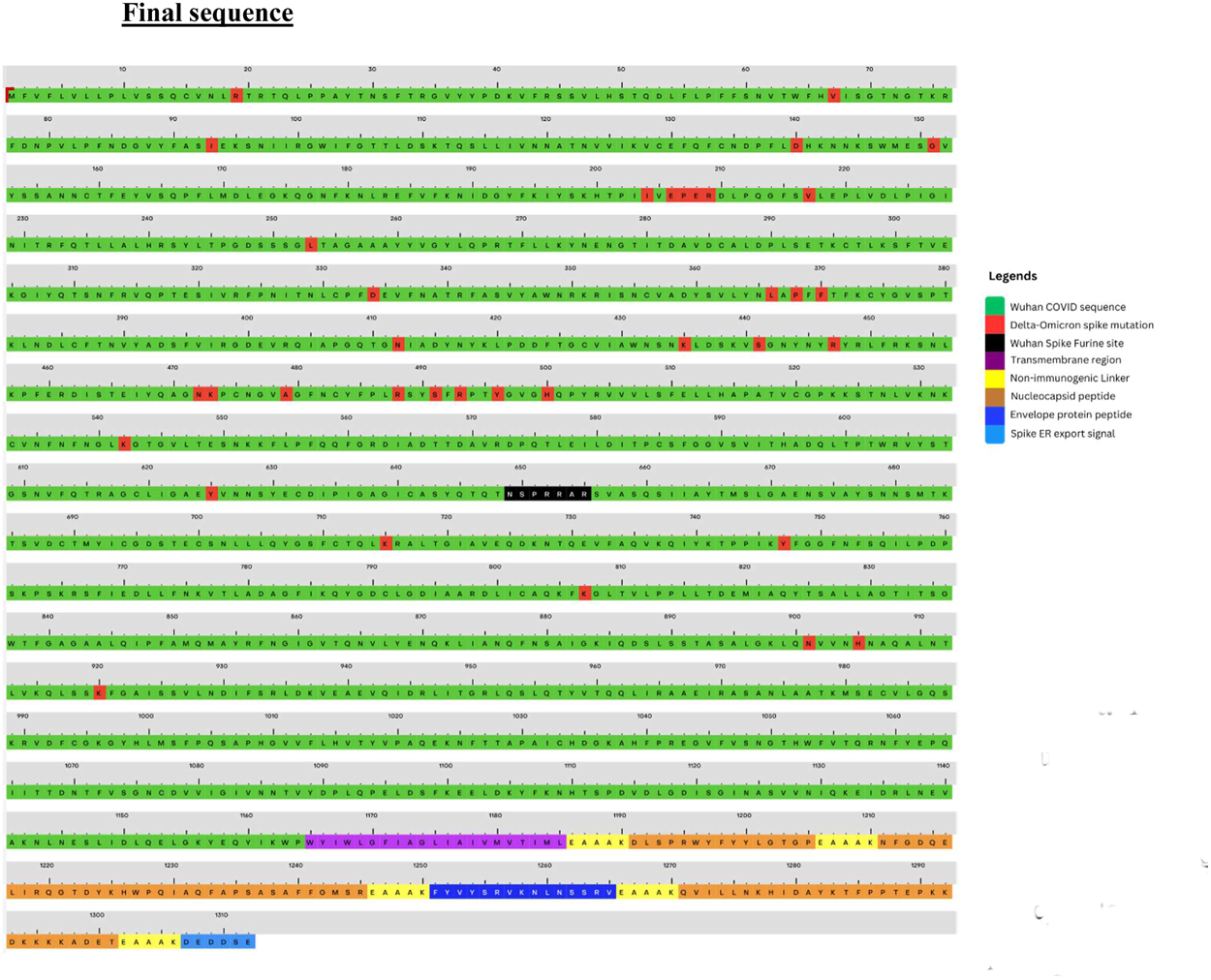
Core β-strand (dark blue, in dotted rectangle, residues: 711-729) is essential for pre- to post-fusion transition of spike. This strand maintains the core β-sheet by exchanging interactions with other two strands from same protomer to homologous strands of different protomer of trimeric spike. A) Pre-fusion spike S2 apparatus comprising central helix, HR region and core β-sheet. B) Post-fusion spike S2 apparatus comprising central helix fused with HR region; Core β-sheet with exchanged 2^nd^ and 3^rd^ strands between protomers. Core β-strand remains unchanged. C) Central helix in pre and post-fusion conformation. [Reprinted with permission from PNAS (43)].

## De-novo vaccine structure modelling

The resulting sequence was imported in to the Robetta server which uses the RoseTTAFold algorithm to build protein structures and using the default settings the vaccine structure modelled. In addition, the model building was also achieved using the AlphaFold (v2.0) in Uppsala server (UPPMAX, Uppsala Multidisciplinary Center for Advanced Computational Science, Uppsala University, Sweden) ^56^. The Pymol (PyMOL-Molecular Graphics System, Open-Source version, Schrödinger, LLC) and the alpha-viewer python package (Severin Dicks, IBSM Freiburg) were used for the analyses and visualization of the structures. The models for hybrid spike without transmembrane domain and wild type SARS-CoV-2 spike were also generated by AlphaFold. Furthermore, all three models were compared using the pTM (predicated template modelling) mode to obtain PAE (predicated aligned error) map. Finally, the predicted local distance difference plots (pLDDT) were calculated for all models and carefully checked the pLDDT values which indicate per residue confidence scores for the accuracy of the models.

## Molecular Dynamics Simulations (MDS) Protocol

### System preparation and Trajectory production

The three-dimensional structure of the final vaccine model (1273 a.a) residues was prepared using the default protocol of PlayMolecule ProteinPrepare ^57^. The protonation states of all residues at pH 7.0 were determined by PROPKA v3.1 ^58^. The missing atoms were added, the bond orders corrected, and the H-networks (atoms) were optimized by PDB2PQR v2.1 ^59^. The structure was then minimized using YASARA minimization server and the resulting structure was used for molecular dynamics simulation ^60^.

The molecular dynamics (MD) simulation of the system was performed for 100 ns using the TIP3P (transferable intermolecular potential 3P) water model as the solvent ^61^. The protein structure was placed in a orthorhombic box with a minimum distance of 10 Å between the protein and the box edges and solvated with water molecules and 0.1 M (molar) NaCl solution. The system was neutralized by adding eight sodium ions to balance the charge of the protein. The temperature and pressure monitored using the NPT ensemble of Berendsen thermostat 300 K and barostat 1 atm for the structure ^62^. The production run was set to 100ns time, and the trajectories recorded using the OPLS2005 force field ^63^. The total number of atoms in the full system checked and noted 311510.

### Simulation analysis

The trajectories were analysed for various structural and dynamical properties, such as root-mean-square deviation (RMSD), root-mean-square fluctuation (RMSF), radius of gyration (Rg), time-based gradients analysis.

## Results

### UniProt analyses

Uniprot mining of SARS-COV-2 structural proteins (spike, nucleocapsid, membrane and envelope protein) were divided into 3 parts: a) Retrieval of SARS-COV-2, SARS, common cold and MERS sequences, b) Clustal Omega alignment of each structural protein from Uniprot and c) Sequence similarity based neighbouring-joining tree generation (without distance values) in Clustal Omega. At first, sequences corresponding to coronaviruses infecting all host animals were considered. For nucleocapsid sequences, we found out inclusion of MERS nucleocapsid generates a distinct tree which slowly departs from other corona including SARS-COV-2 (Fig 2A). Further, we generated circular tree by filtering out gapped sequences, protein fragments and non-human host infecting corona viruses. This confirmed that MERS nucleocapsid protein sequences indeed form a distinct tree, but by being more conserved as compared to others (Fig 2B). In fact, it did not even change much within the MERS family (Fig. 2B). Usually, nucleocapsid protein is highly conserved than spike and more immunogenic. It is also essential for maintaining RNA structure, packaging inside viral particle. If random mutations arise in nucleocapsid, virus will not survive in long run. Taking this MERS scenario, introduction of recurrent random mutations in nucleocapsid protein (by immunological perspective) may be beneficial for vaccine design. Although, SARS-COV-2 (SARS-2) nucleocapsid has more mutations than MERS, still the frequency is much low compared to spike (Fig. 2B, Supplementary Fig. S1). Alongside we also generated similar sequence-similarity trees with other structural proteins (Spike, Membrane and Envelope) (Supplementary Fig. S1, S2 and S3). The focus was on 2 aspects: 1) Conserved structural proteins among living corona viruses (MERS, SARS-COV-2 and common cold corona viruses; not SARS as it is already extinct by natural immunity); 2) How immunogenic these proteins are. There we found out that envelope protein also can be a suitable candidate like nucleocapsid. These protein sequences were highly conserved among SARS-COV-2 strains (Supplementary Fig. S3). Thus, we have taken one non-allergenic epitope from envelope protein which is important for binding to human cell junction proteins and has robust immune response. Surprisingly, spike protein sequences were also found highly conserved among MERS (Supplementary Fig. S1). As already chosen in existing vaccine candidates including our design, not much study was done on this further. Membrane protein (M) was found highly conserved among extinct SARS strains (Supplementary Fig. S2) and immunologically less persistent (described below), therefore not chosen.

### Literature hunting for potential T-cell epitopes for vaccine design

Immunogenic sequences of COVID nucleocapsid protein and envelope protein were carefully shortlisted from literature studies. Initially, the attention was on the sequences with overlapping CD4+ and CD8+ epitopes. Similarity with common cold viral sequences were also taken into consideration. The data available from recovered patient and asymptomatic population were given highest importance. Final chosen peptides were screened based on their structures (from PDBsum) or from secondary structure predicted by Phyre2 server (in absence of PDB for envelope protein; Supplementary Fig. S7). As known, SARS-2 CD8+ epitopes are hotspots of nonsynonymous mutations to cause short-lived immunity. Since CD8+ epitopes are mostly unstructured, special consideration was made to choose epitopes with definite structures to attract CD4 based immunity. Subsequently, two epitopes were found folded in Alpha Fold. For the rest, we found one epitope fully and another partially non-structural after validation in Alpha Fold (described below). They were kept in the final construct based on their ability to provoke instantaneous immune response due to high similarity with common cold and strong memory response. Membrane protein epitopes were found to be less persistent as per the literature study with exception of highly COVID-exposed hospital workers who are more immune to COVID than any other population. The chosen peptides are described in Supplemental Table 1.

### RoseTTAFold building and structure validation in AlphaFold

The designed sequence had signal peptide, transmembrane domain and a cytoplasmic domain formed by T-cell epitopes interconnected by EAAAK linker. Major stabilizing portion of putative ADCC/ADE region and core β-strand were removed. The resulting local disorder in remaining ADE segment after deletion was confirmed from analysis in AlphaFold-disorder package (Supplementary Fig. S11) which can predict residue fluctuations from pLDDT values and relative solvent accessibility (RSA) calculated. The positively charged Endoplasmic reticulum (ER) exit signal from COVID native spike was placed at the cytoplasmic end to facilitate easy exit from ER. This sequence was first modelled in RoseTTAFold and found to be structurally similar to native spike all domains including cytoplasmic domain are distinctly well formed (Fig. 7). To validate further, we have modelled in Alpha Fold for two sequences: with and without transmembrane (ΔTM) region. It was seen the at least two epitopes (NFGDQELIRQGTDYKHWPQIAQFAPSASAFFGMSR and FYVYSRVKNLNSSRV) were folded in ΔTM sequence (Fig. 8). While first sequence is well folded in the TM+ (with transmembrane) construct, but not the other (Fig. 10). It appears the first sequence is the major driver of folding for the designed cytoplasmic domain. Also, inter-residual distance analysis in PyMOL showed longer distance (21 Å) between this peptide and last sequence or memory epitope (QVILLNKHIDAYKTFPPTEPKKDKKKKADET) in TM+ construct than the other (Fig. 9). The other peptide (DLSPRWYFYYLGTGP) and memory epitope are fully unstructured in TM+ construct. Memory epitope is partially structured in ΔTM possibly due to closer distance (8.5 Å) (Fig. 11A) to folding driver peptide. Seemingly, this closeness is resulted from absence of transmembrane domain in ΔTM which otherwise pulled away (Fig. 9B) the folding driver epitope from memory epitope in TM+ model. This might have caused unstructured memory peptide in TM+ construct. In fact, both TM+ model and native spike (having transmembrane region) have shown mostly unstructured cytoplasmic domain which is further validated by PAE scores similarities (Supplementary Fig. S9, S10) in this domain for these two models. For the main protein, the receptor binding doman (RBD) is found to be well folded and identical among all three models including native spike (Supplementary Fig. S4). Also, the core β-strand is replaced by two discontinuous stretches of β-strands which can form hydrogen bonds with the complementary strand in the core β-sheet (Supplementary Fig. S5). Thereby, the structural integrity of this region can be maintained. Although, the lengths of these strands are not enough to support pre- to post fusion transition of spike unlike native core β-strand.

**Fig 7:**
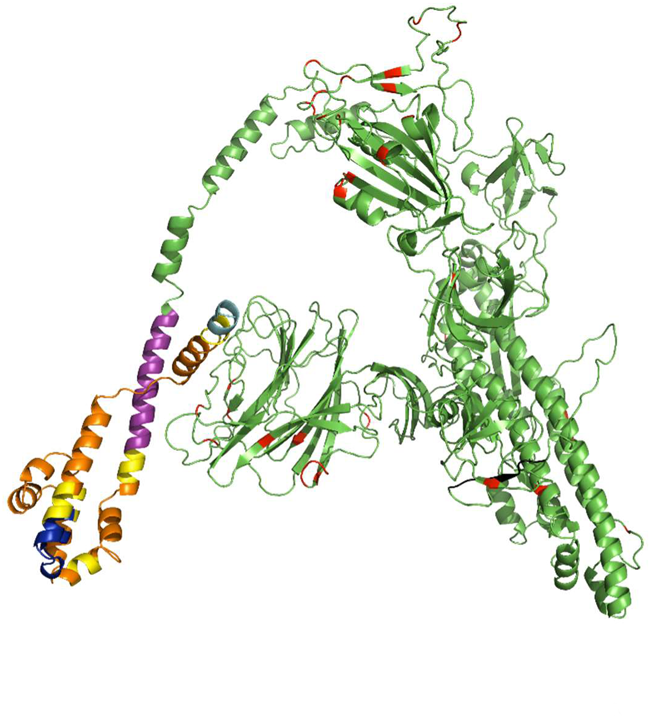
Sequence and 3-D model (RoseTTAFold) of the TM+ vaccine construct. The mutations corresponding to Delta and Omicron variants are coloured red. Transmembrane domain, linkers, cytoplasmic T-epitopes and terminal ER exit signals are coloured differently. All colour codes are explained in the legend on right side. The T-epitopes appear helical in this model. Overall, the cytoplasmic domain looks like well folded coiled coil located close to the main protein.

**Fig 8:**
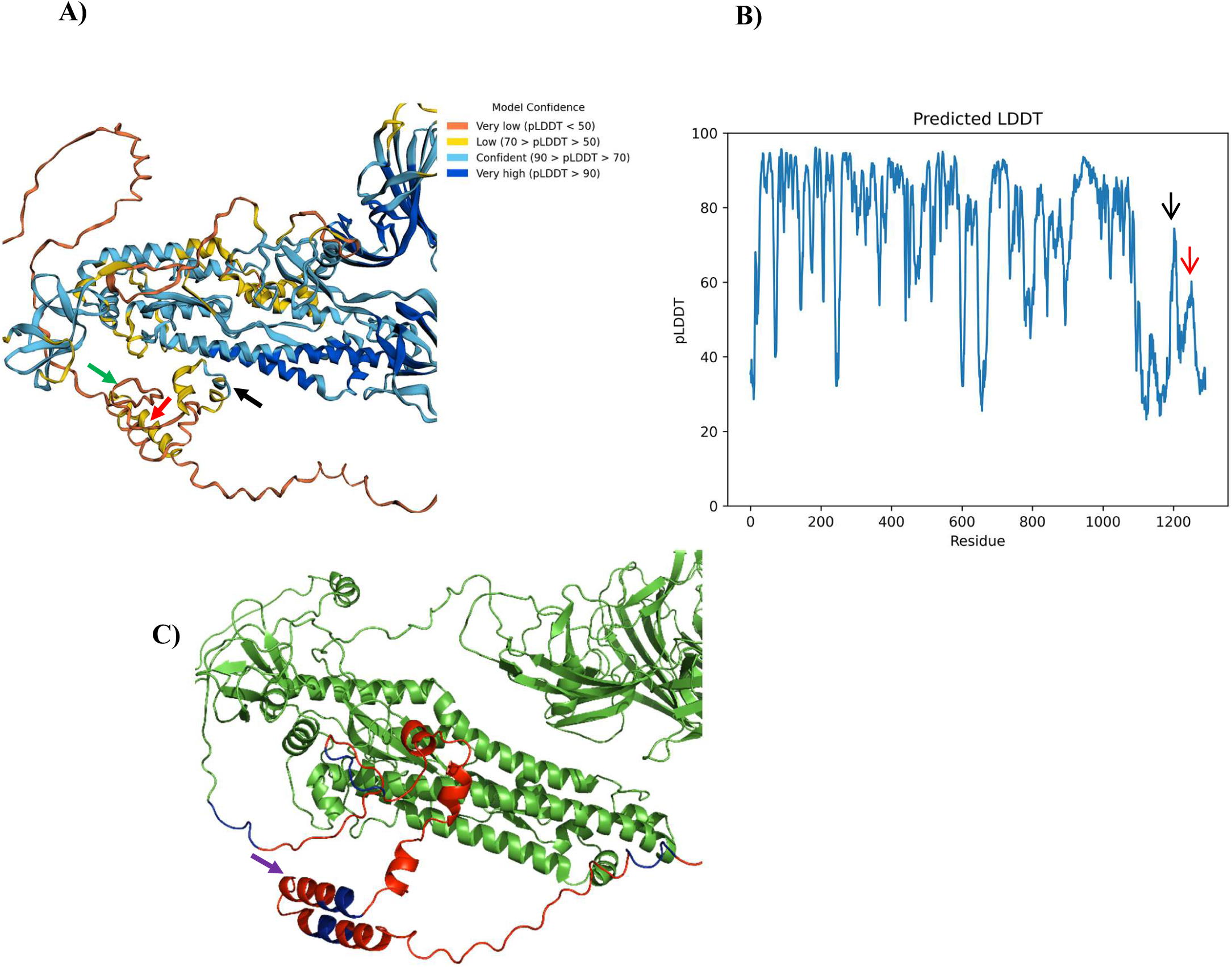
AlphaFold model of the final vaccine construct (without transmembrane domain) appears as well folded. **A)** Best model for the ΔTM construct and **B)** Predicted local distance difference test (pLDDT) plot. Except cytoplasmic domain, most of the other portions of the molecule have pLDDT values >70. The model is colour coded according to model confidence/pLDDT values in B. In cytoplasmic domain, the central region of folding driver epitope (marked by black arrow) has pLDDT value >70 and >50 for terminal portions. Similarly, part of memory peptide (red arrow marked) was folded with >50 pLDDT value. As expected from noncomplementary β-stranded structure in PDBsum, the first epitope (DLSPRWYFYYLGTGP) (green arrow marked) forms unstructured region (pLDDT<50). **C)** The envelope peptide between folding initiator peptide and memory epitope is folded as helices (from PyMOL; Purple arrow marked). The cytoplasmic epitopes and EAAAK linkers are coloured as red and blue respectively. Rest of the protein is coloured as green.

**Fig 9:**
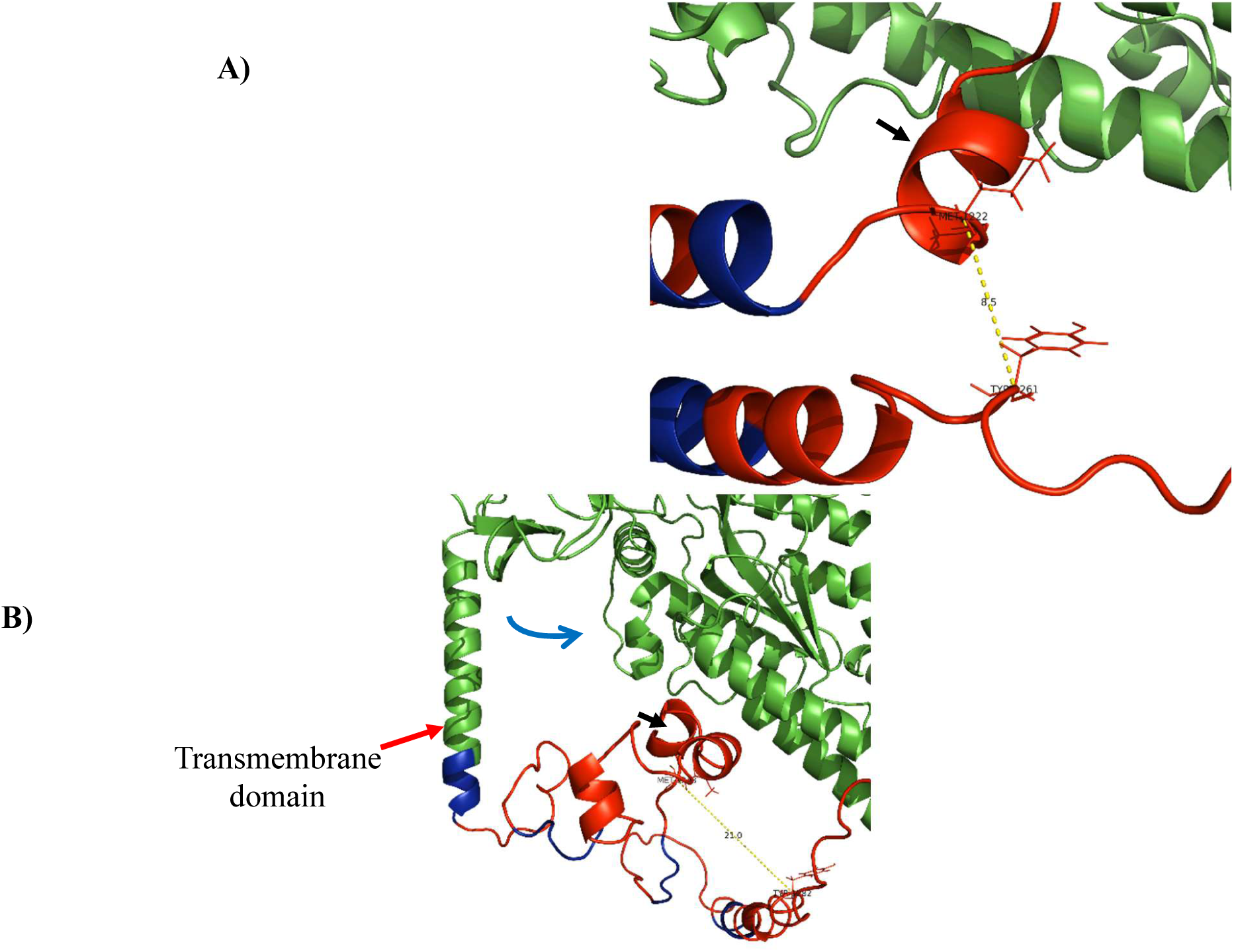
Folding centre of the vaccine constructs with T cell epitopes (red) joined by linkers (blue). **A)** Without transmembrane domain (ΔTM) and **B)** with transmembrane domain (+TM). The linkers (EAAAK) are coloured as blue while individual T-epitopes in cytoplasmic domain are coloured as red. The closest distance between folding driver epitope (FDP) (marked by black arrow) and memory peptide is monitored by central methionine of the former and a tyrosine residue of the latter. In ΔTM, this distance is 8.5 Å which widens to 21 Å in +TM construct. The transmembrane domain (marked by red arrow) is helical and seems to pull (blue arrow) the adjacent cytoplasmic domain regions (which includes folding driver epitope) towards rest of the protein (spike portion, green). The distal region (with memory peptide) of cytoplasmic domain are not influenced by this pulling effect and stays away from FDP.

**Fig 10:**
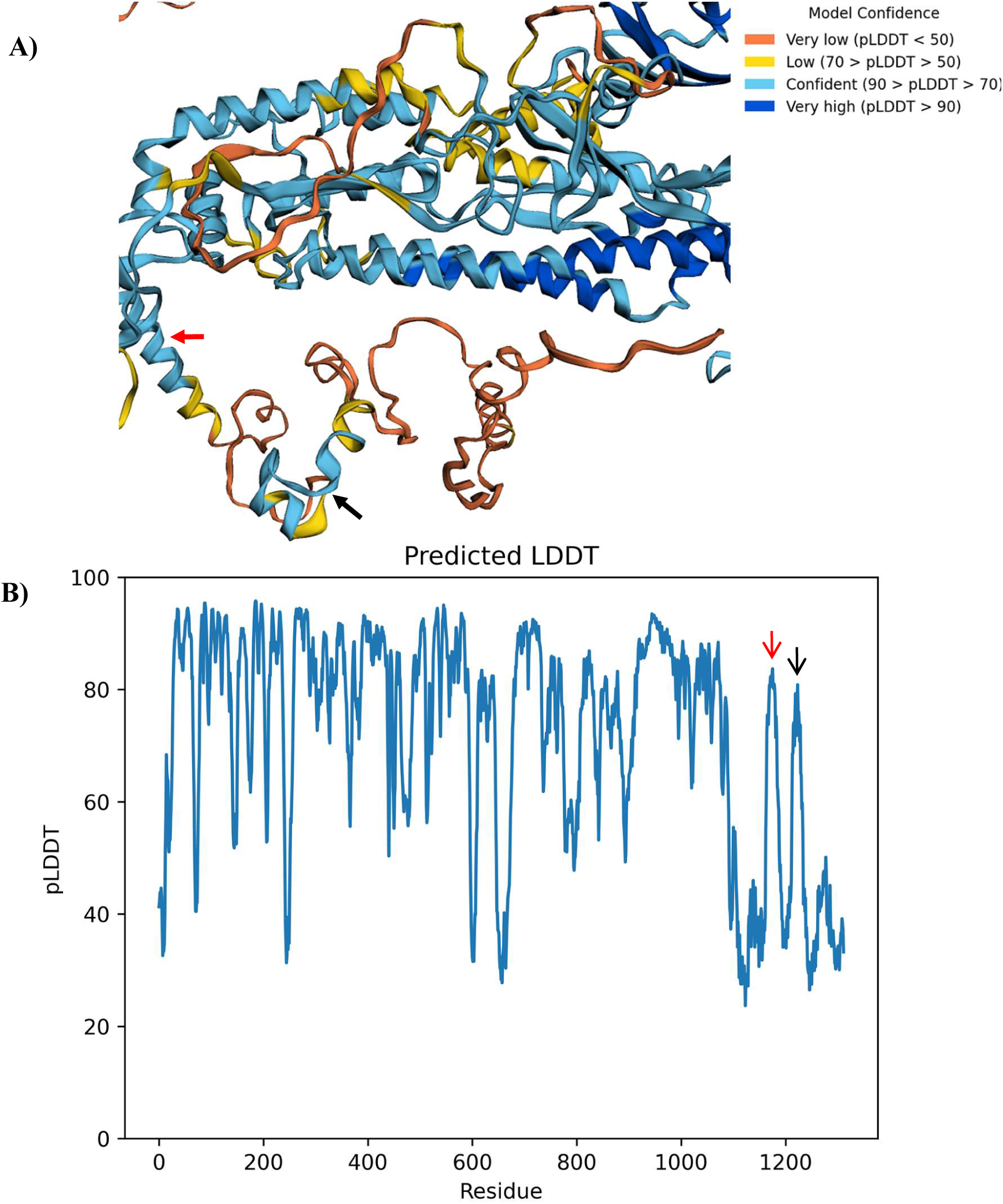
AlphaFold model of the vaccine construct with transmembrane domain (+TM) and predicted LDDT plot. **A)** The best model is shown. The folding initiator epitope is marked by black arrow. This epitope is well folded. Other epitopes are unstructured in cytoplasmic region. The transmembrane domain is marked by red arrow. **B)** The pLDDT plot shows that the red arrow marked transmembrane region (with pLDDT>80) and most of the other domains are well folded except cytoplasmic region. The model confidence for folding driver peptide (denoted by black arrow) is higher (≈80) than ΔTM indicating no effect from neighbouring unstructured epitopes.

**Fig 11A:**
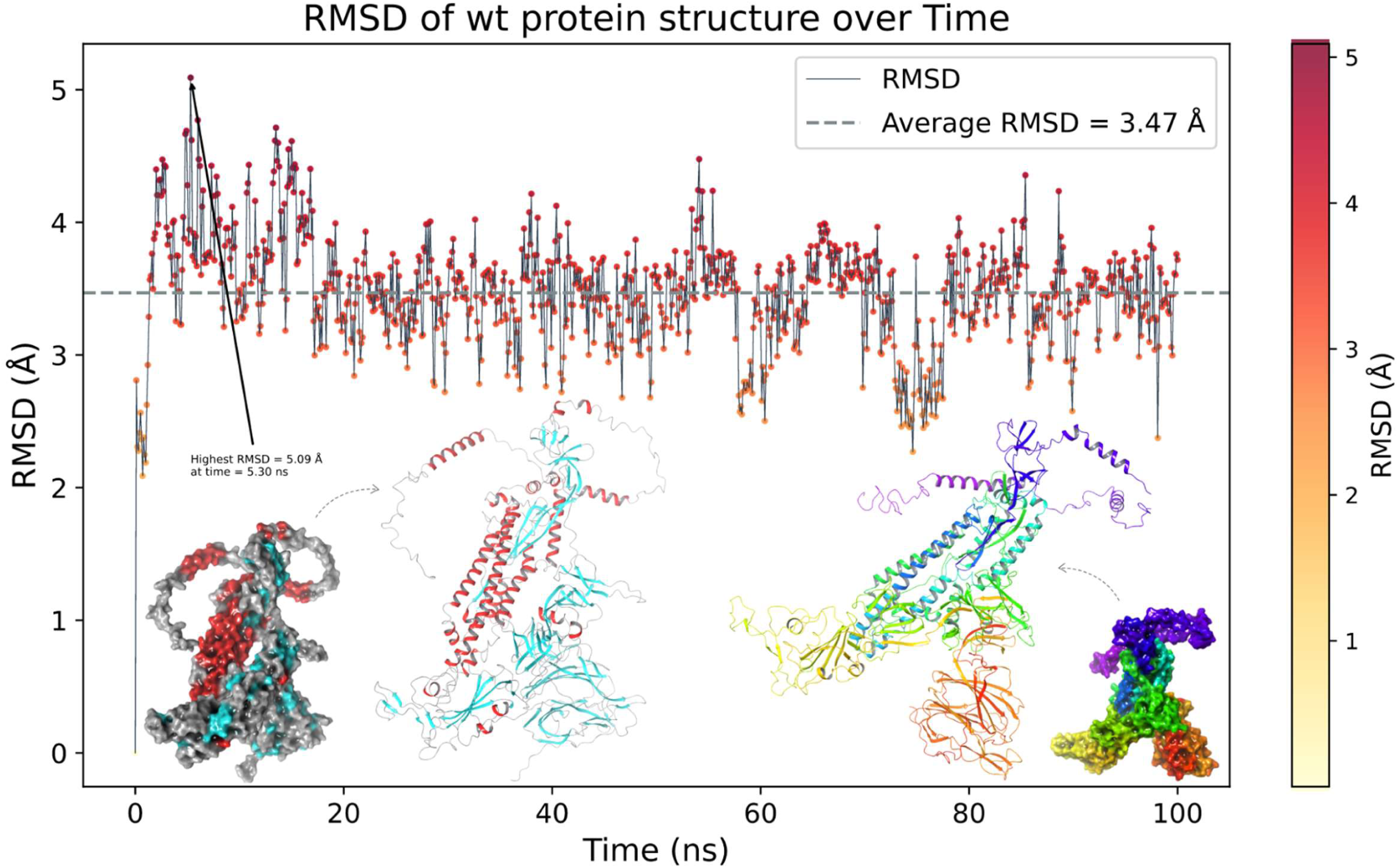
Shows the highest RMSD (root mean square deviation) at 0ns (5.09 Å) and average 3.47Å for overall structure of the C-alpha group atoms. The average line is shown in grey color.

**Fig 11B:**
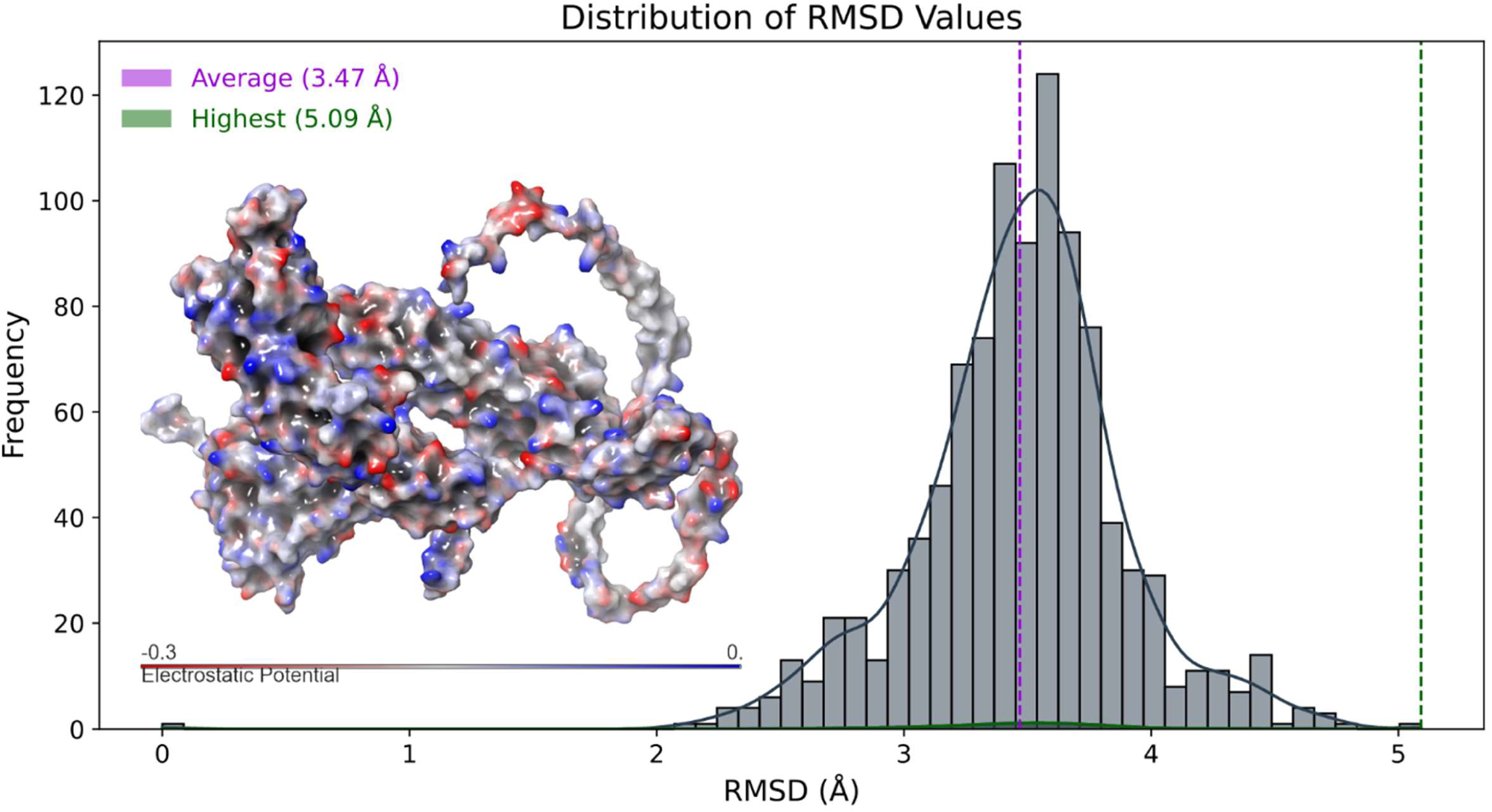
Shows the highest RMSD (root mean square deviation) at 0ns(5.09 Å) and average 3.47Å for overall structure of the c-alpha group atoms including the electrostatic interactions value of -0.3.

### Design Validation through Molecular Dynamics Simulations

One of the challenges of de novo design of anti-variant COVID-19 vaccines is to ensure that they elicit robust and long-lasting T-cell memory response. In this study we further validated the AlphaFold designs in a 100ns of molecular dynamics (MD) simulations and analyzed various parameters of root mean square deviation (RMSD), root mean square fluctuation (RMSF), and radius of gyration (Rg) and assessed the structural stability and dynamics of the design. These parameters can provide insights into the conformational changes, flexibility, and compactness of the structures over time and indicating stable design. Figure 11A shows the root-mean-square deviation (RMSD) plot of the protein backbone atoms as a function of time. The RMSD is a measure of how much the protein structure deviates from a reference structure, usually the initial or native structure. The initial RMSD value at 5.30 ns was 5.09 Å, indicating a large deviation from the starting structure. This could be due to the initial equilibration of the system and the relaxation of the protein conformation. After that, the RMSD values decreased and fluctuated around an average value of 3.47 Å, suggesting that the protein reached a stable conformation and did not undergo significant structural changes during the rest of the simulation. The protein structure is determined by a balance of different forces, including covalent bonds, van der Waals interactions, hydrogen bonds, salt bridges and electrostatic interactions. Electrostatic interactions are caused by the presence of charged or polar residues on the protein surface or in the interior. Electrostatic interactions can affect protein structure in several ways. They can stabilize or destabilize the protein fold by forming favourable or unfavourable interactions with other residues or with the solvent molecules. They can also modulate the flexibility and dynamics of the protein by influencing the rigidity or mobility of certain regions or domains. The histogram in figure 11B shows the electrostatic interaction values between -0.3 for the protein structure at 100ns time. The electrostatic interaction value is calculated as the sum of Coulombic interactions between all pairs of charged or polar atoms in the protein, divided by the number of atoms. A more negative value indicates the overall attractive, while a positive value indicates an overall repulsive. The histogram shows that most of the electrostatic interaction values are close to zero, indicating that there is a balance between attractive and repulsive interactions in the protein structure. However, there are some outliers with more negative values, indicating that these regions or residues have somewhat stronger electrostatic interactions than others. These residues with more negative values may be involved in stabilizing salt bridges or H-bonds with other residues and region and they could be more rigid or flexible than others, affecting their conformational changes or dynamics at 100ns of time trajectory. The root-mean-square fluctuation (RMSF) analysis is a method to measure the flexibility of different regions of a molecular structure over time. It can reveal the more stable or high dynamic residues or regions of the structure contributing during molecular dynamics simulations. Figure 12A shows the RMSF values of each residue along the protein sequence. The residues with higher RMSF values are coloured in red, indicating that they are more flexible than the residues with lower RMSF values, which are coloured in grey. The most flexible residue is 1246-, with an RMSF value of 14.48 Å and is located at the C-terminus of the protein, which is expected to be more mobile than the rest of the structure residues. In addition, the C-terminal region, starting from 1160-THR to the end, has an overall high RMSF value, suggesting that it is very flexible region. Also, average RMSF value of 3.69 Å for overall structure in 100ns time of simulations. This can also be seen in the histogram in Figure 12B, which shows the distribution of RMSF values indicating the significance of the c-terminal region in terms of flexibility and conformational transitions. The radius of gyration (Rg) is a measure of how the atoms of a protein are distributed around its centre of mass. The smaller the Rg, the more compact the protein structure is. In figure 13A-B, the Rg of the protein was calculated for each frame of a 100ns molecular dynamics (MD) simulation. The highest value was 20.59 Å and the average value was 20.33 Å, indicating that the protein maintained a relatively compact conformation throughout the simulation. To visualize the structural changes over time, we plotted the time-based gradients of the structures in figure 14, where the blue colour represents the initial structure, and the red colour represents the final structure. The structure underwent shrinkage during the simulation, which could be due to the formation or stabilization of intra- or inter-molecular interactions. The three-dimensional representation of the protein structure at different time points during the simulation can be seen in Figure 14. The structures are aligned by their centres of mass and coloured according to their time order. The blue structure corresponds to the initial conformation at 0 ns, while the red structure corresponds to the final conformation at 100 ns. The intermediate structures are coloured with a gradient from blue to red, indicating their relative position in time.

**Fig 12A:**
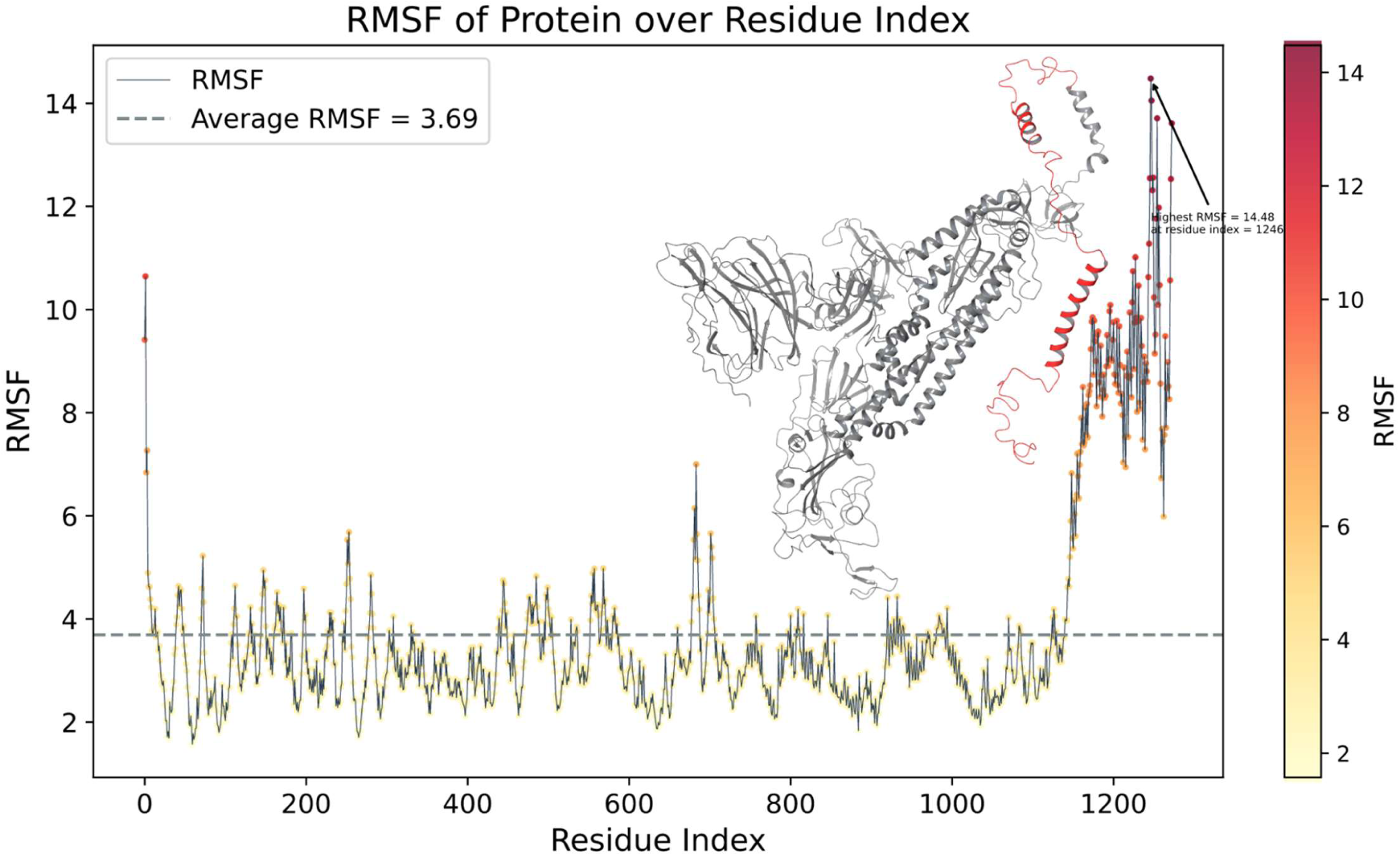
Shows the average 3.69 Å RMSF (root mean square fluctuation) and highest for single reside at 1246 position 14.48 Å at 100 ns time. The highly flexible region is represented in red color.

**Fig 12B:**
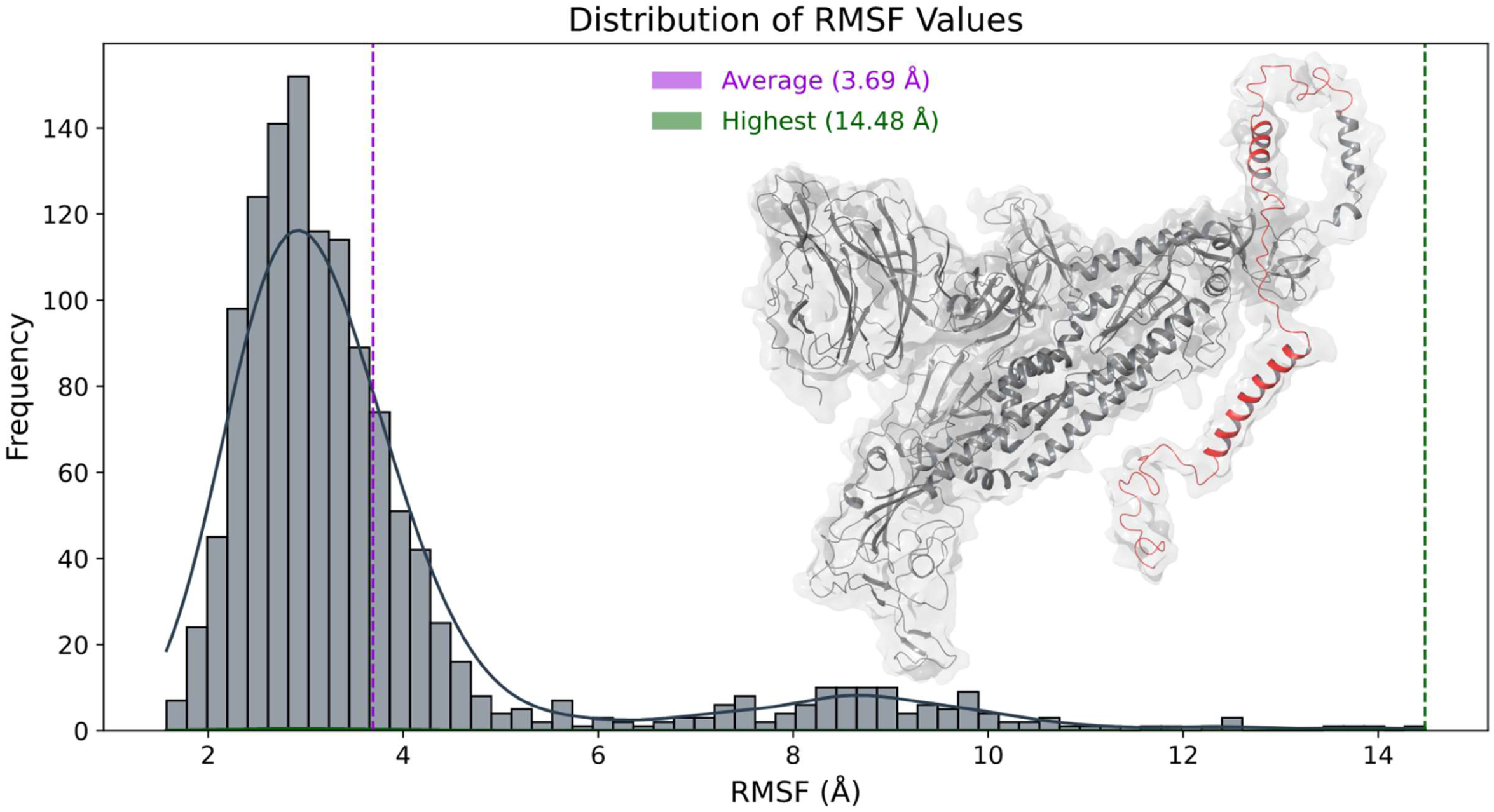
Shows the RMSF values in a histogram representation and the average value of 3.69 Å in pink line color while the green indicates the highest value of 14.48 Å at 100ns trajectory.

**Fig 13A:**
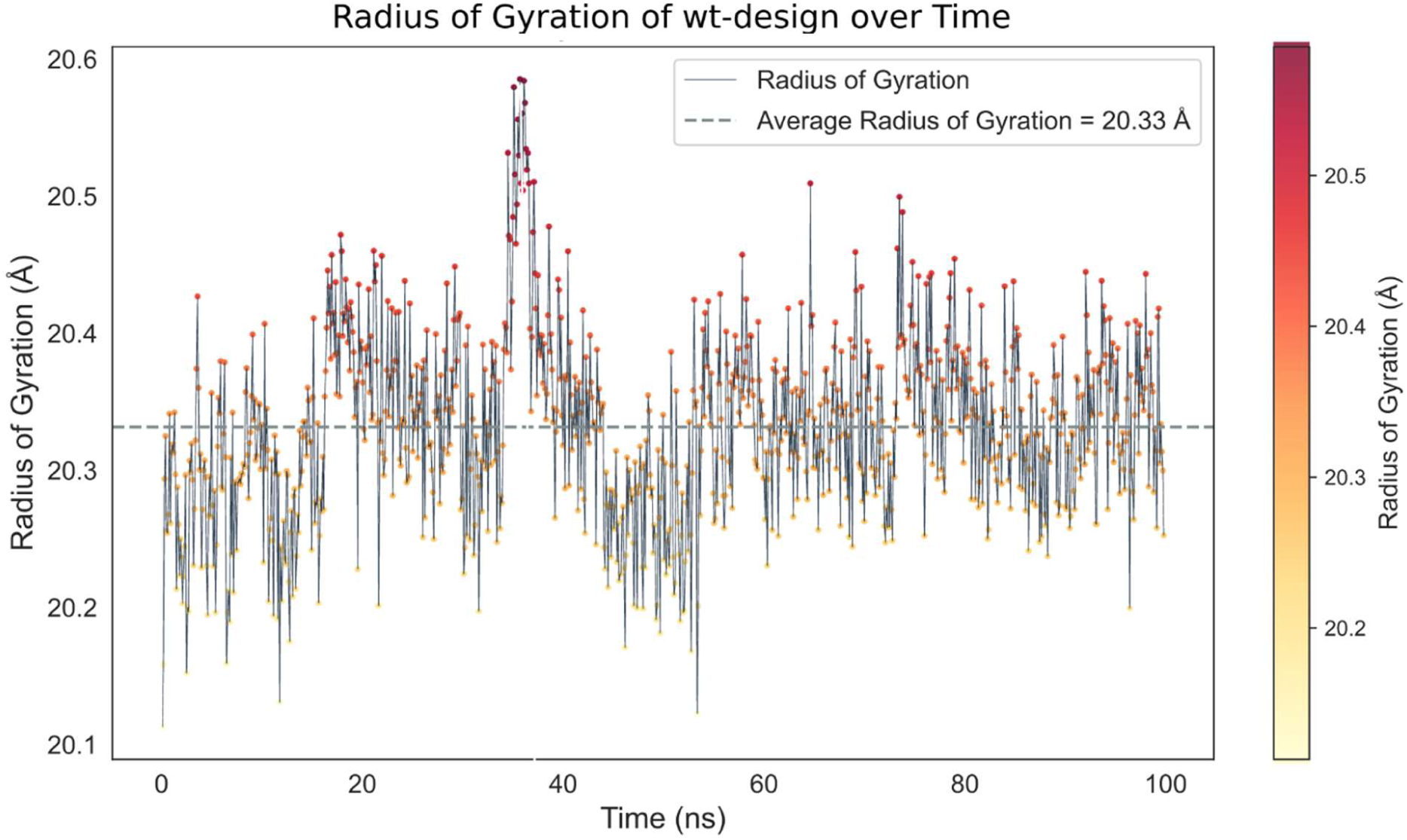
Indicates the Rg(radius of gyration) for the structure at 100ns time simulations with average value of 20.33 Å in a grey color line representation.

**Fig 13B:**
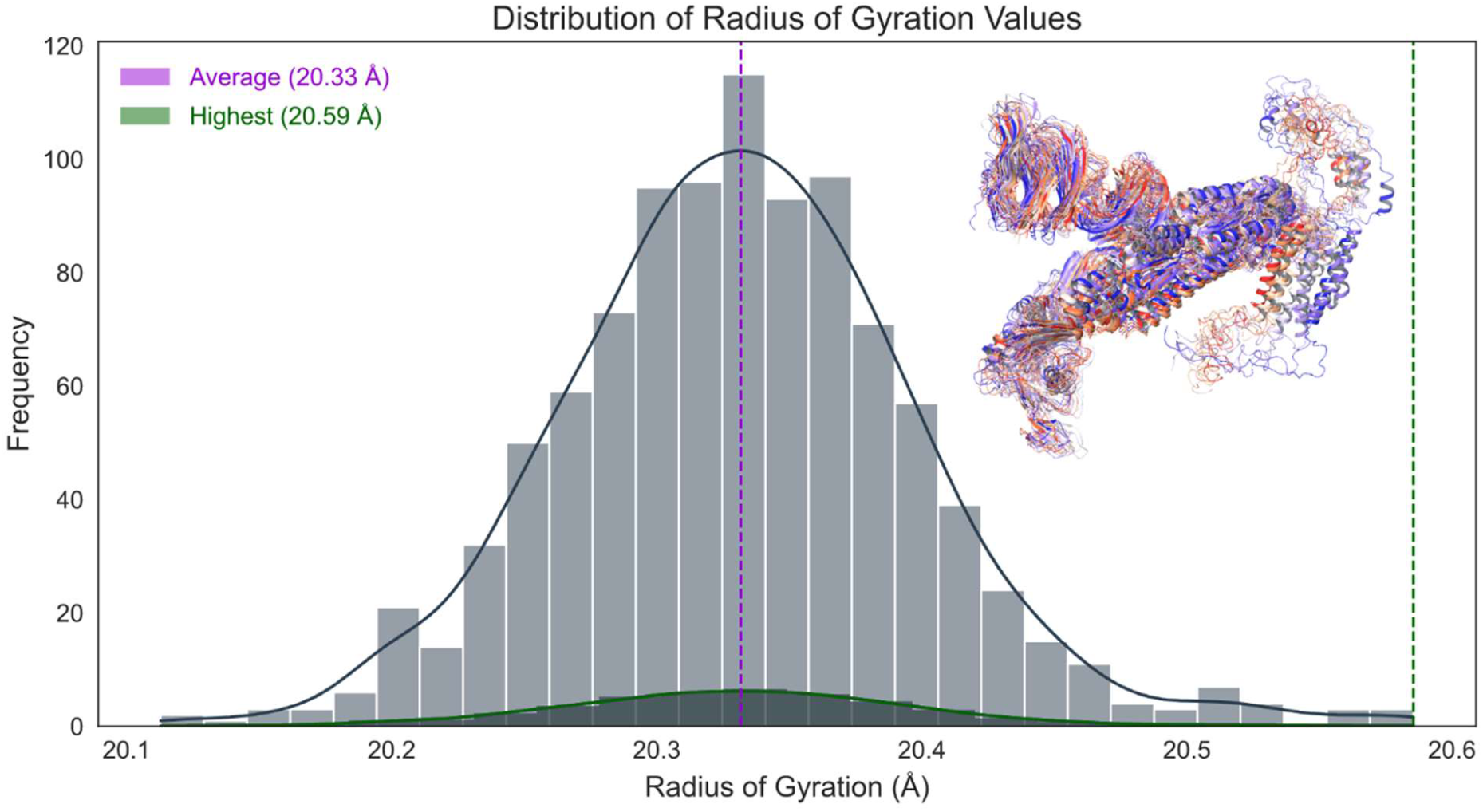
The histogram shows the Rg values of highest 20.59Å including average 20.33 Å in pink and green color respectively

**Fig 14.**
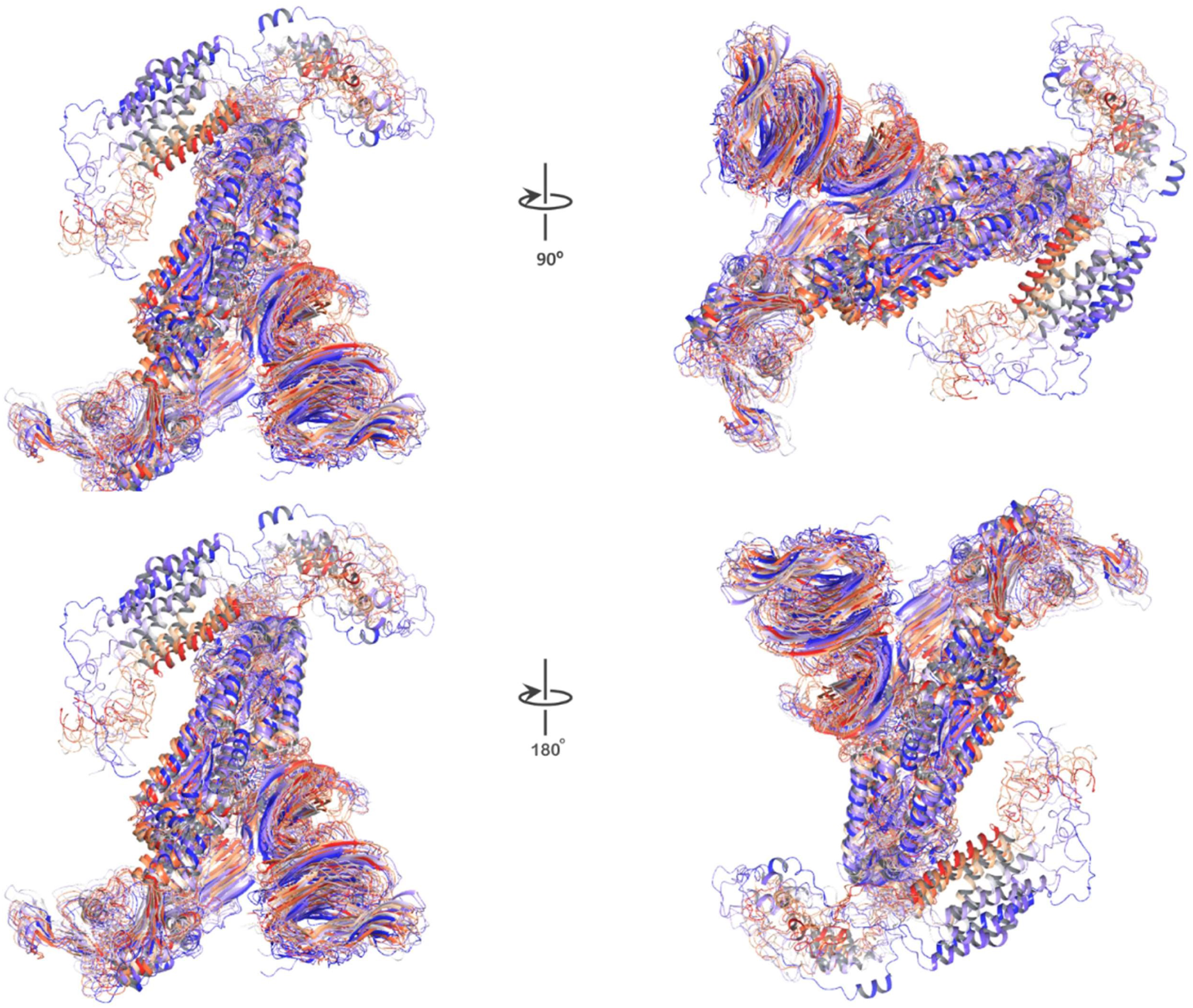
shows the design structures change over time from 0ns to 100ns in two different angles: 90° and 180°. The blue color indicates the initial state, and the red color indicates the final conformation and compactness that the structures achieve after 100ns of simulation.

## Discussion

In this study we applied computational design and simulation of two vaccine structures with potential to overcome SARS-CoV-2. The other conserved structural proteins along with spike were targeted for these potential vaccine constructs. These structures were further validated via 100 ns molecular dynamics simulations. Our vaccine constructs are designed based on SARS-CoV-2 structural proteins’ T-cell evoking epitopes screened from literature. The hallmark of current COVID19 vaccination is wild type spike protein because of its potent antigenicity. But the presence of vaccine evading variants and mutations led us to incorporate the mutant spike in our design. For e.g. the deadly delta variant from pre-vaccination era was followed by milder omicron variants after mass vaccination. This massive paradigm shift in viral strains predicted future variants will be more likely recombinants to evade vaccine elicited immunity. Therefore, hybrid spike sequence with both delta and omicron mutations is important for the potential design of vaccine candidates which we achieved in our design. Furthermore, spike protein’s S2 segment has regions which do not generate neutralizing antibodies and do not have dominant T cell response. There is also a short cytoplasmic domain. Our designs replaced it by inserting COVID’s T cell epitopes interconnected by non-immunogenic, helical & rigid linker sequence EAAAK. To incorporate all these linkers, we also have removed segments of core β-strand (buried deep inside spike core, structurally dispensable in pre-fusion spike and immunologically less significant) (Fig. 4, Fig. 6) from the S2 region of spike. This can significantly be useful in the design to add a new segment to the full molecule without making it larger in size and conformation to reduce the cost and time of synthesizability. Additionally, the absence of this region can prevent the spike to go from pre- to post fusion conformation (Fig. 6). Thus, standard proline-proline (non-viral sequence) insertion which is used in other spike-based vaccines is not considered including the putative region in S1 which caused ADE responses in SARS as per literature study and structural analysis (Fig. 5B). The selection of T-epitopes to build cytoplasmic domain, non-allergenic peptides among nucleocapsid’s numerous T cell epitopes (Fig. 3) were enlisted including memory peptide which helps in eliciting strong immune responses in recovered and vaccinated patients. These sequences are also highly similar to other common cold coronavirus (HuCoV) nucleocapsid protein sequences (Fig 1). As all humans have immune response against HuCoV from childhood driven by pulmonary immune memory including CD4+ epitopes. Interestingly, using these epitopes has an advantage to generate long term and quick immune response. This establishes their importance in persistent immunity. Whether the selected peptides cause misfolding in the generated construct or not, the PDBsum analysis helped to screen nucleocapsid peptides which were found to fold with correct secondary structures (Fig 3B). For envelope peptide without any PDB structure, we have relied on Phyre2 result (Supplementary Fig. S7). Later model building in RosettaFold showed these peptides as alpha-helical in the designed cytoplasmic domain of the construct. Final Alpha Fold model building predicted that at least two peptides are folded while the other two epitopes are mostly unstructured in construct without transmembrane region. One of the folded epitopes can be the main player in folding for other subsequent peptides. The predicted LDDT values were >70 for this epitope in both ΔTM and +TM sequences (Fig. 8, Fig. 10). Also, we found out that the transmembrane domain pulls the adjacent region including the abovementioned principal epitope for folding in cytoplasmic domain from the other portions in +TM construct. The resulting distance between last epitope and folding driver epitope is much more in +TM construct than ΔTM construct. Interestingly, the folding driver peptide was constructed by joining numerous CD4+ and CD8+ responsive N-peptides from immunological studies (Supplemental Table 1). In native Nucleocapsid protein, they are located as flanked to each other, parsed by immune system as numerous small peptides. Apart from immunological significance, they might play a role in folding of native nucleocapsid around viral RNA. Parallel molecular dynamics in 100ns simulation time revealed stable conformations of the design structures in terms of RMSD, initially higher and then onwards much more stable and very flexible C-terminal region in the RMSF. Furthermore, Rg and time based-gradient analysis revealed a compact system suggesting overall conformational stability of the design structures and less mobility or fluctuation transitions except highly flexible C-terminal regions (which otherwise will be fragmented by immune systems once recognized as potential epitopes).

## Conclusion

Given the utmost need for potential vaccines against deadly SARS-CoV-2 pathogen candidate recently an emergency left by WHO, our study provides comprehensive in-silico strategy to design vaccine against SARS-CoV-2 using the long-lasting potent T-cell immune memory strategy. Our strategy proposes that the ΔTM construct with linker joined cytoplasmic epitopes can be a potential vaccine construct against the deadly SARS-CoV-2 (VOC) including extending its capability further against the future evolving variants which is further validated in a 100ns simulations revealing the stable overall conformations and less structure mobility or fluctuation transitions. Hence this strategy, not only for SARS-CoV-2 (VOC) but can easily be replicated for other viruses with potential to cause pandemic such as Monkey pox, influenza, respiratory syncytial, dengue and other pox related viruses. Our study provides new insights and a comprehensive in silco protocol for long-lasting potent T-cell immune memory by considering a significant set of mutations from SARS-CoV-2 (VOCs) in the vaccine constructs. Furthermore, in-vivo, and in-vitro validation of the vaccine construct and other analysis are necessary which is lacking in this study including the incorporation of hotspot residues of spike protein of other variants and variants of concern to design a pan coronavirus vaccine.

## Authors contribution

AG and MMG planned the study and vaccine designs. AG performed for the alignments, AlphaFold analysis, RoseTTAFold and finalizing epitopes for the vaccine design. MKS processed sequences to obtain AlphaFold models. SU performed and led the molecular dynamics simulations (MDS) and designed the Graphical Abstract of the study. AG, SU and MMG wrote the paper.

## Supporting information

Supplementary Fig S1 (Enlarged)

## Acknowledgements

The Alpha Fold modelling was enabled by resources in project [SNIC 2022/22-709] provided by the Swedish National Infrastructure for Computing (SNIC) at UPPMAX, partially funded by the Swedish Research Council through grant agreement no. 2018-05973.

## Authors information

AG: Protein Crystallographer and Bioinformatician

MKS: Protein and organic Crystallographer, post-doctoral Scientist in Uppsala University, Sweden

SU: Structural Biologist

MMG: Scientist G (Retired), ICMR-National Institute of Virology, India (Deceased)

