## Supplementary figures and images for "*De novo* design of anti-variant COVID-19 Vaccine"

### Supplementary Fig S1 (Enlarged)

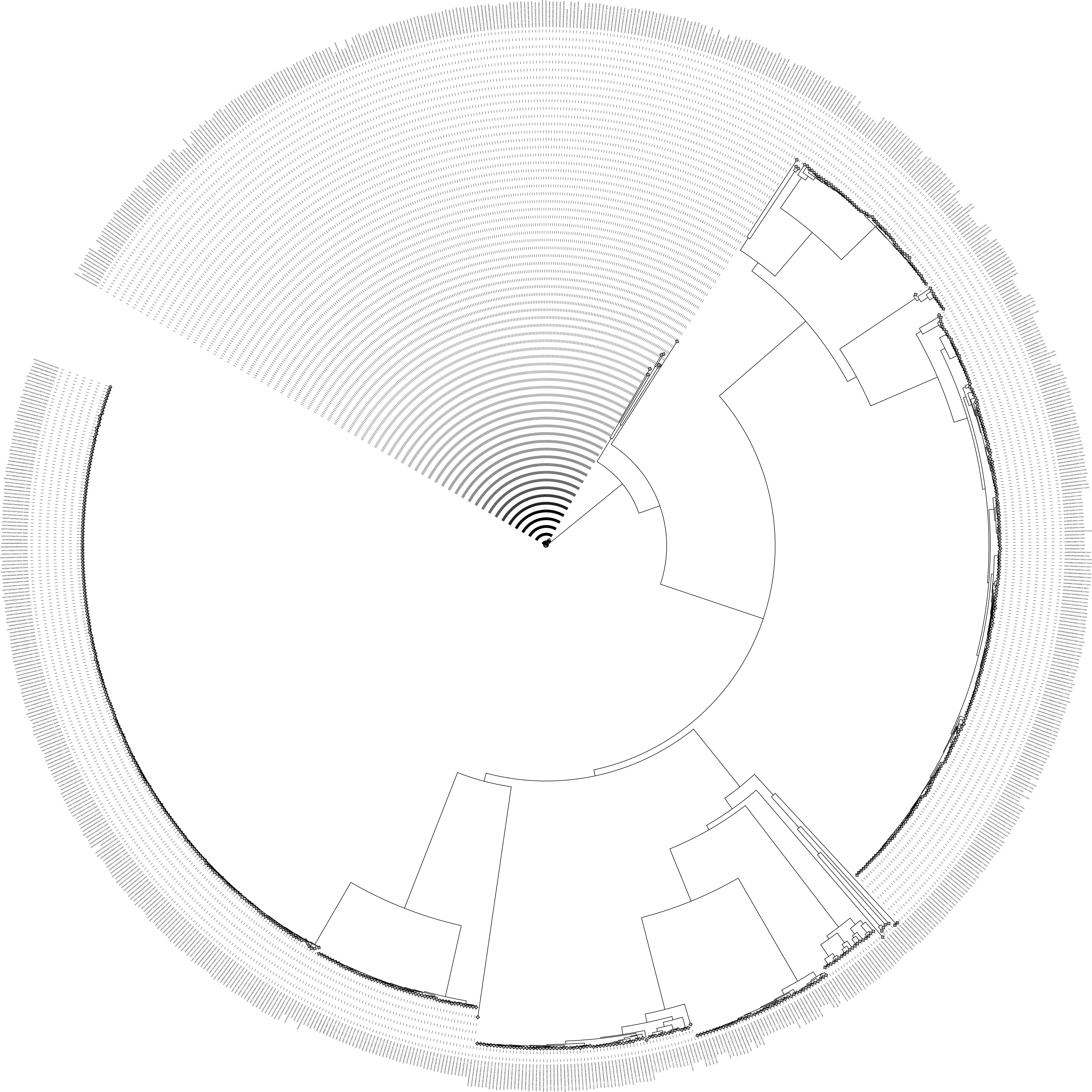
