## Supplementary Fig. S 1-11 for "*De novo* design of anti-variant COVID-19 Vaccine"

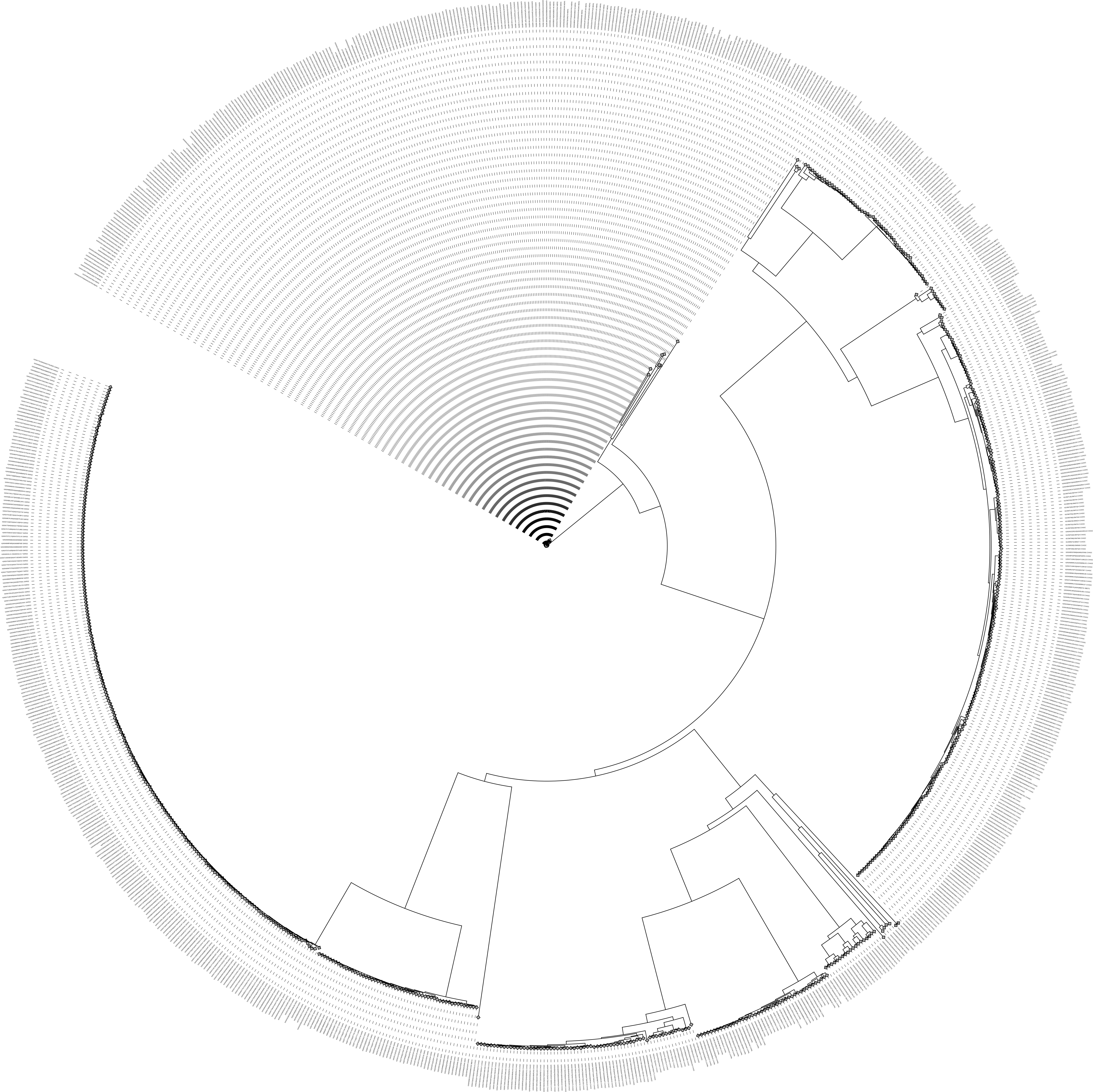
**Fig S1: Sequence similarity based** **neighbouring-joining circular tree of spike proteins from SARS, SARS-2, MERS and common cold coronaviruses.** Marked by arrow are MERS spike protein family (chosen from Uniprot after removing fragments, gapped sequences and non-human host infecting MERS coronaviruses) showing very little changes in their sequences as compared to others.

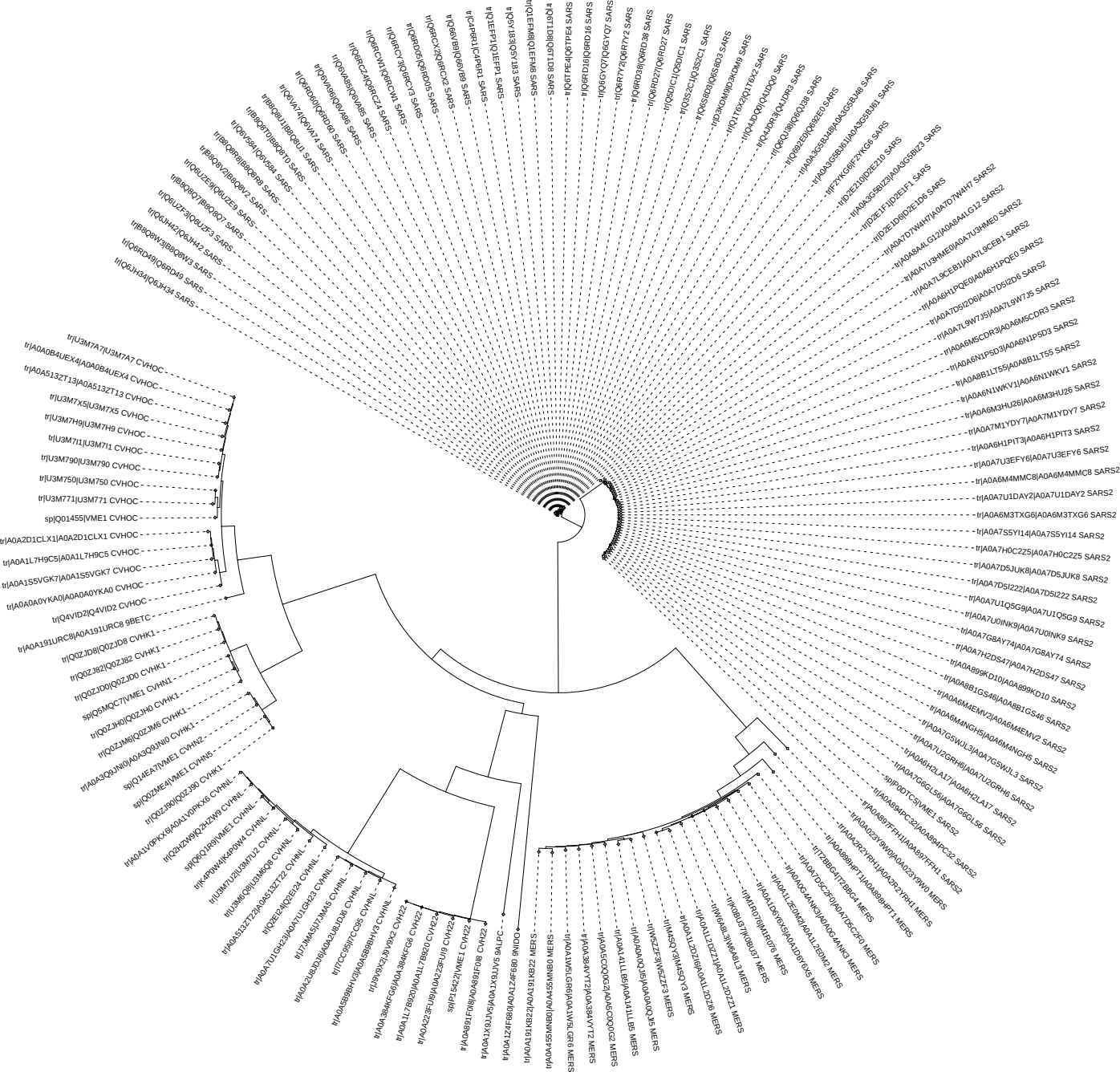

**Fig S2: Sequence similarity based** **neighbouring-joining circular tree of membrane proteins from SARS, SARS-2, MERS and common cold coronaviruses.** Indicated by arrow are SARS membrane protein family (chosen from Uniprot after removing fragments, gapped sequences and non-human host infecting coronaviruses) showing very little changes in their sequences as compared to others.

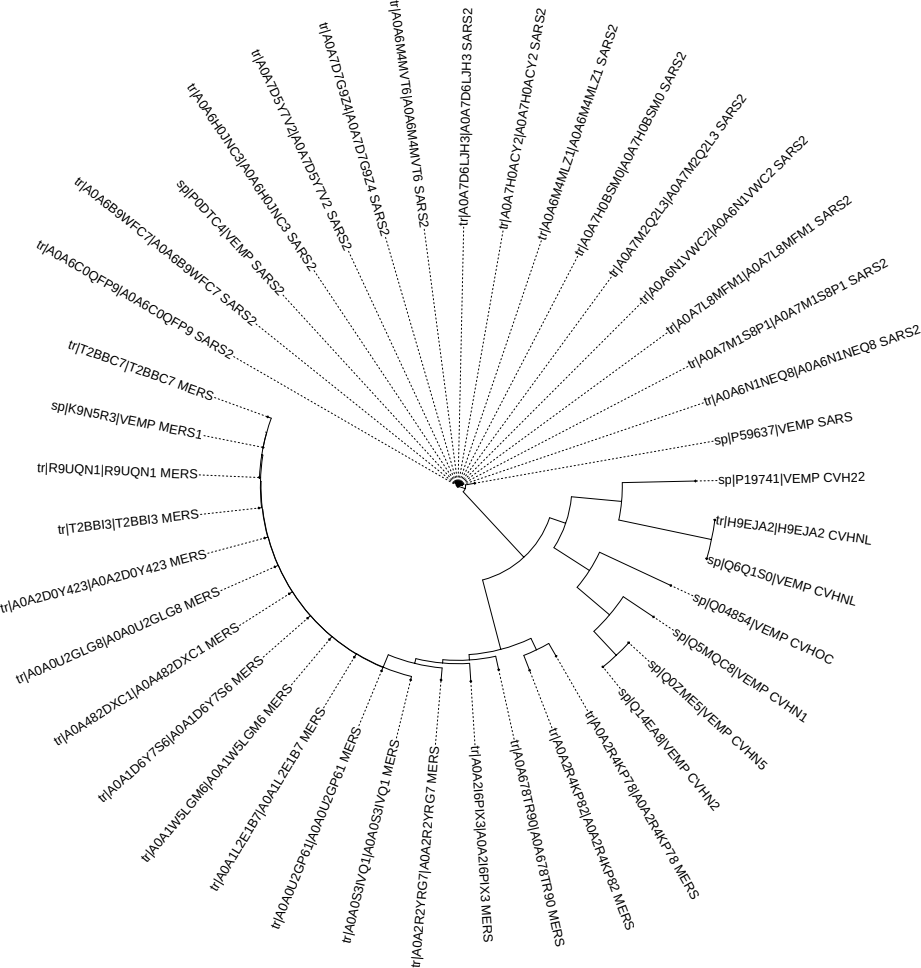
**Fig S3: Sequence similarity based** **neighbouring-joining circular tree of envelope proteins from SARS, SARS-2, MERS and common cold coronaviruses.** Marked by arrow are SARS-2 (Covid) envelope protein family (chosen from Uniprot after removing fragments, gapped sequences and non-human host infecting coronaviruses) showing very little changes in their sequences as compared to others. For SARS, only one sequence was found to be without gaps.

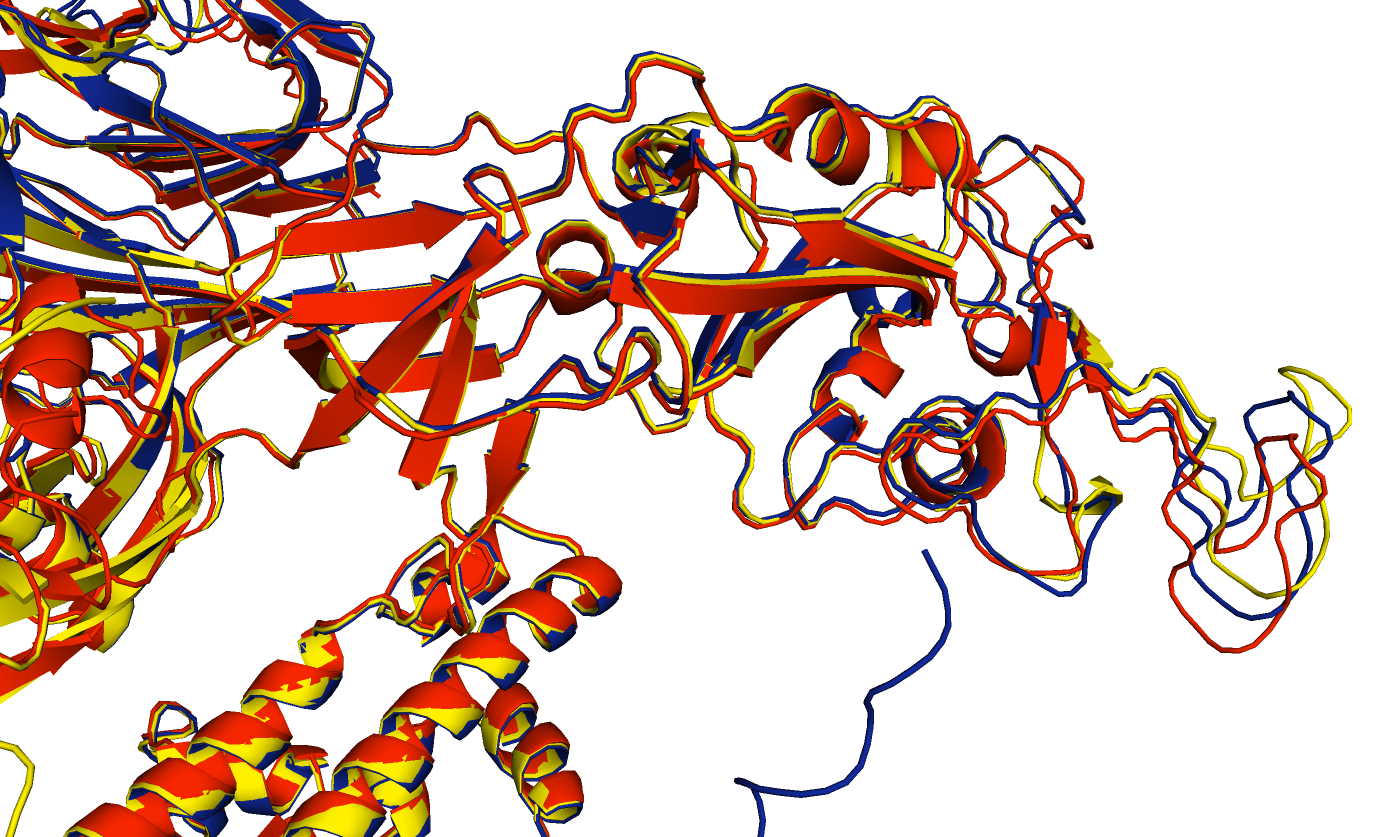
**Fig S4: Receptor binding domains of +TM, ∆TM and Native spike models appear nearly identical.** Vaccine models with (yellow) and without transmembrane (blue) domain were aligned on native spike model (red) in PyMOL. Encircled are RBDs of all three showing very little changes in their structures.

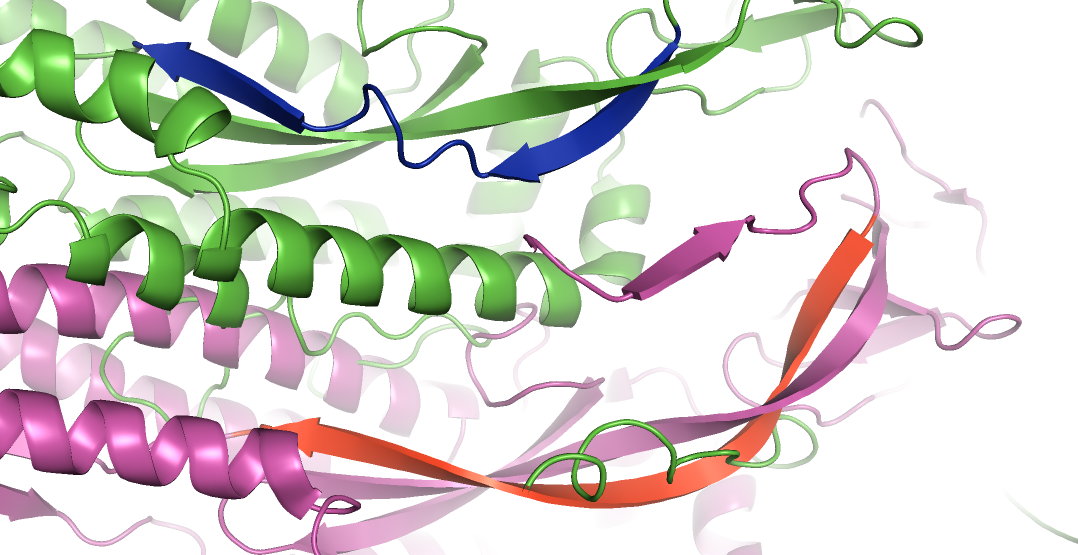

**Fig S5: Native core β-strand is replaced by two discontinuous β-strands in vaccine model.** The long core β-strand (red) in native spike was deleted in vaccine construct (green) and compared with native spike (magenta). It was observed the lost strand is replaced by two adjacent β-strand stretches (blue) which can support the core β-sheet in that region. Thus the structural integrity is maintained. But the novel strands’ lengths will not be sufficient for interactions with other subunits of trimeric spike to facilitate pre- to post-fusion transition.

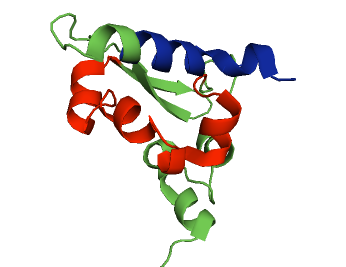

**Fig S6: PDB structure of Nucleocapsid protein shows the interaction between epitopes chosen.** The pivotal epitope for folding (red) and part of memory epitope (blue) are shown in the structure (6ZCO.pdb). The interacting region is marked by arrow.

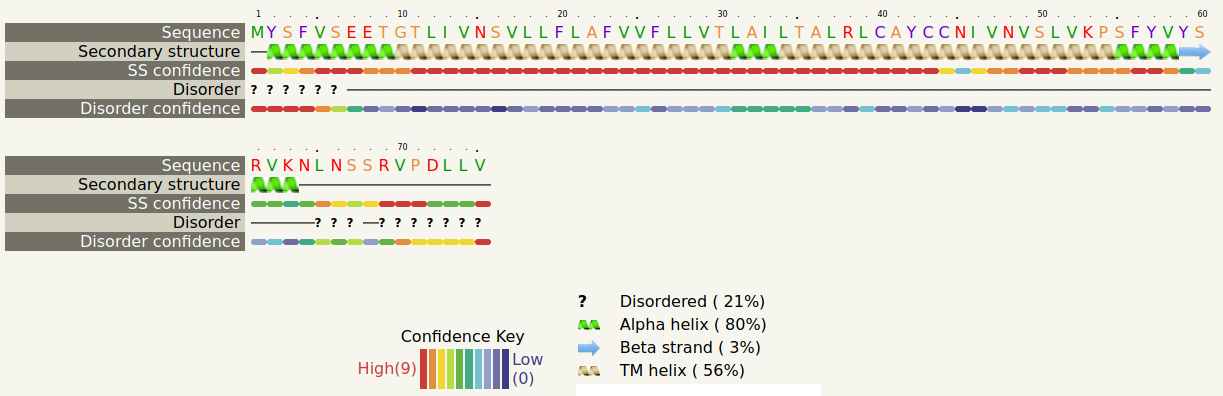

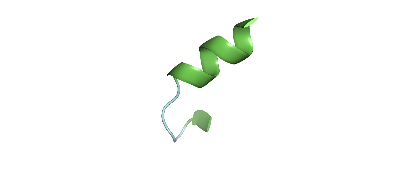

**Fig S7: The secondary structure of Envelope protein.** Secondary structure motifs of envelope protein are displayed from Phyre2 server. The different secondary structures in whole protein and their percentages are explained in the legend. α-helices are green; β-strands are cyan; transmembrane regions are grey and disordered regions are coloured other than these three shades. Also colour bars represent the confidence values for the different motifs. The chosen epitope peptide in vaccine is marked by dotted box. Two α-helices connected by a short β-strand and a C-terminal disordered region form this peptide. After model building, this epitope was folded with two α-helices interconnected by a short loop in ∆TM (Inset box on top of legend). There was no folding in case of +TM.

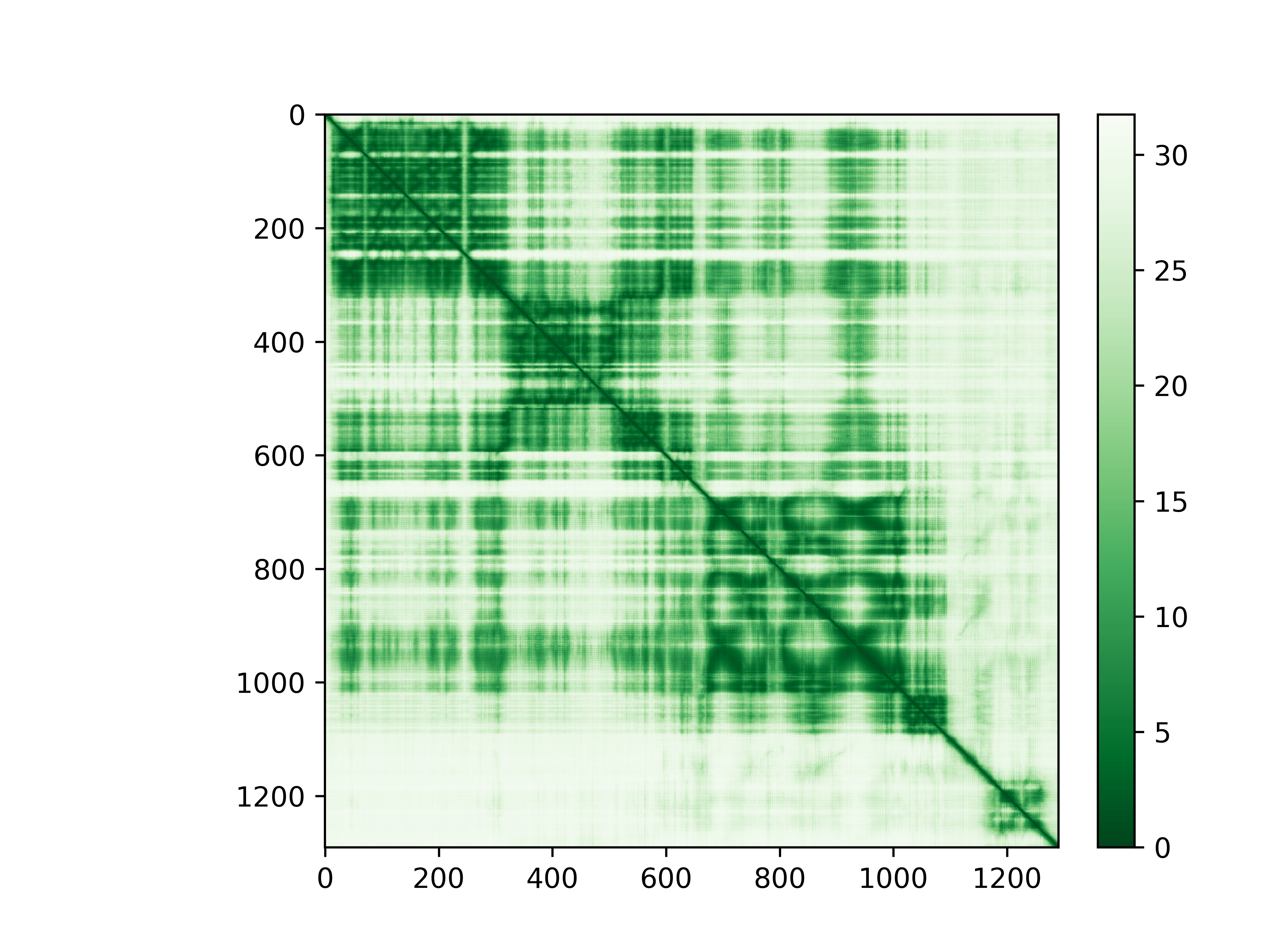

Aligned Residues

Scored Residues

**Fig S8: Predicated aligned error** **(PAE) plot for ∆TM model.** This plot is obtained from monomer pTM (predicated template modelling) method in AlphaFold. Scored residues represent predicted positions of x^th^ residue and aligned residue denote actual positions of y^th^ residue. White gaps indicates maximum positional error (Å). The Accuracy in domain modelling is directly proportional to intensity in green shade in the plot. The colour bar on the right side represents the error levels. Two perpendicular dashed rectangles (red) are drawn on the map which intersects on the diagonal to form a square. Enclosed inside this is the cytoplasmic domain (1165-1291) which is partially disordered**.** Folding driver peptide (1190-1224) is folded with less error (marked by arrow). Adjacent linker and envelope peptide (1230-1244) is also folded with good error values. Towards the end of cytoplasmic domain error increases which indicates unstructured terminus.

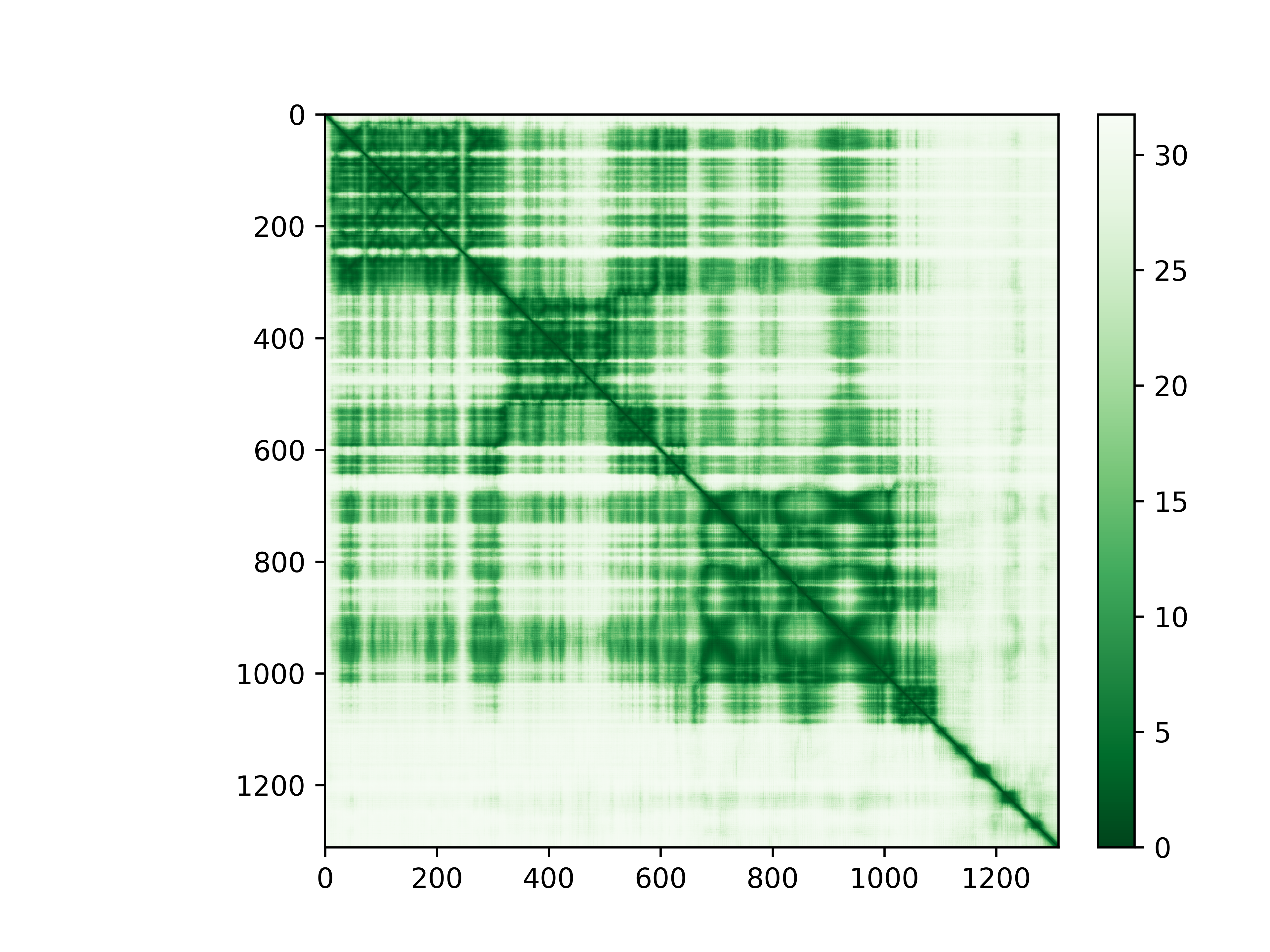

Aligned Residues

Scored Residues

**Fig S9: Predicated aligned error** (**PAE) plot for +TM model.** Heat map is generated from monomer pTM (predicated template modelling) generated model for TM+ in AlphaFold. Enclosed inside two criss-crossed dashed rectangles-generated square box (red) is the cytoplasmic domain (1186-1312). The colour bar on the right side represents the error levels. Folding initiator peptide (1211-1245) is folded with less error (intense green colour). Also this epitope is represented by narrow patch (marked by arrow) on the diagonal as compared to broad and diffused area for ∆TM model in previous figure. This may indicate less effects from neighbouring unstructured regions on this peptide. Overall, in most of the cytoplasmic domain error increases corresponding to disordered structure.

**A)	B)**

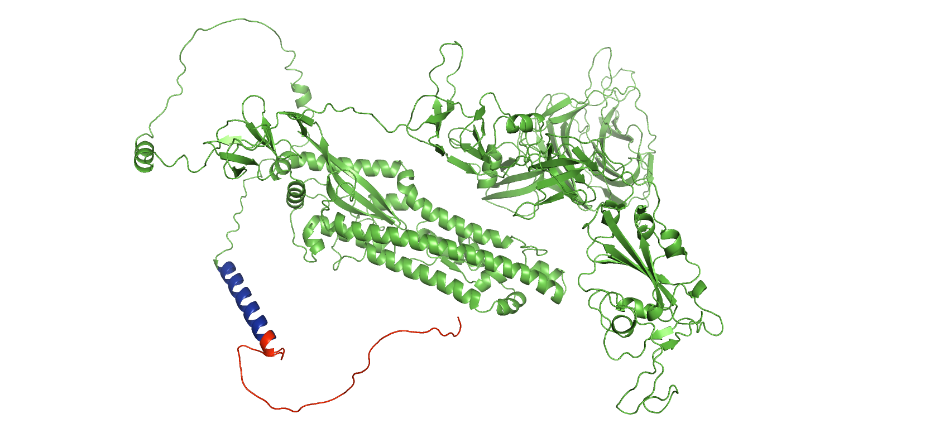

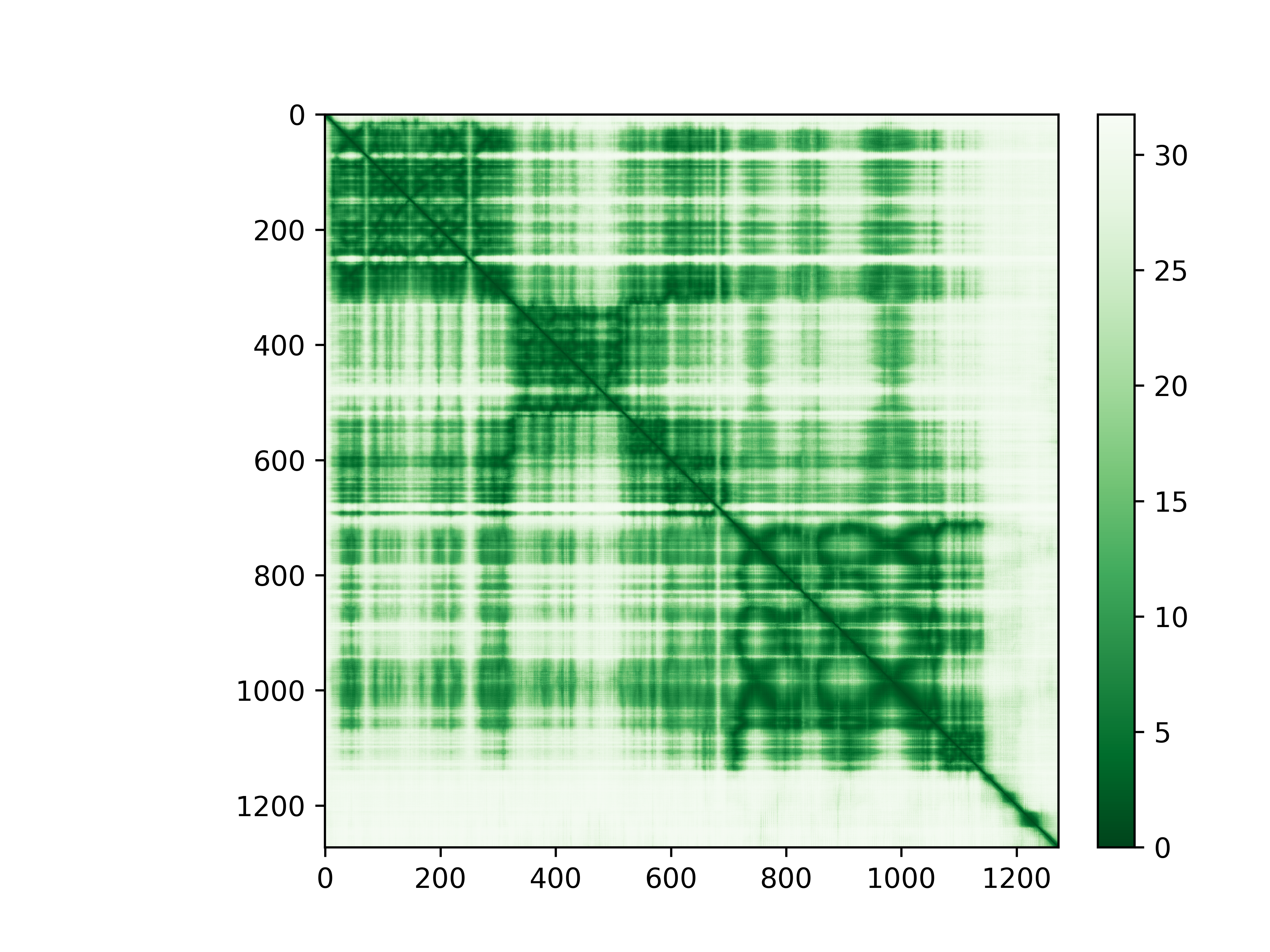

Aligned Residues

Scored Residues

**Fig S10: Predicated aligned error** (**PAE) plot for native spike model. A)** The heat map represents relative errors of domain folding for native spike in pTM model building. Enclosed inside the tiny square (denoted by arrow) generated by two intersecting rectangles (red) is the cytoplasmic domain (1235-1273). The colour bar on the right side represents the error levels. In the cytoplasmic domain white shade dominates corresponding to disordered structure. **B)** Native spike model (best model from AlphaFold visualized in PyMOL) with mostly unstructured cytoplasmic domain (red) and α-helical transmembrane domain (blue). Rest of the protein is coloured as green.

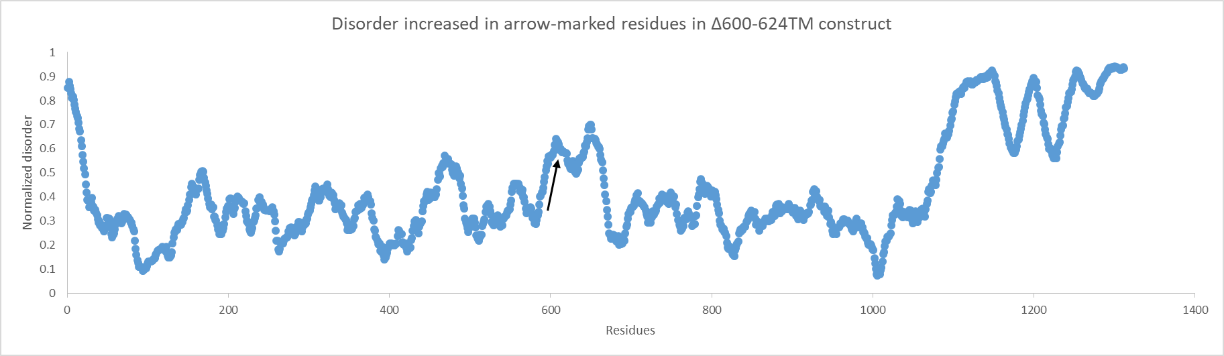
**A)**

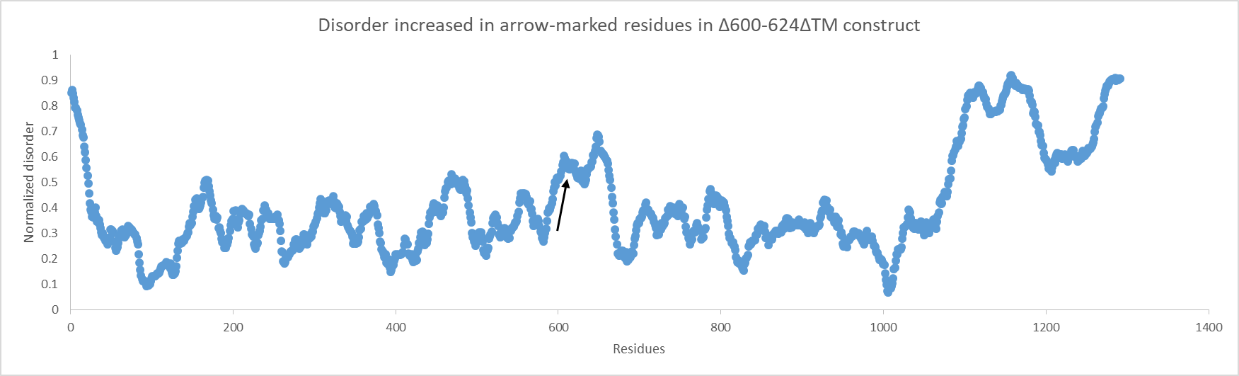
**B)**

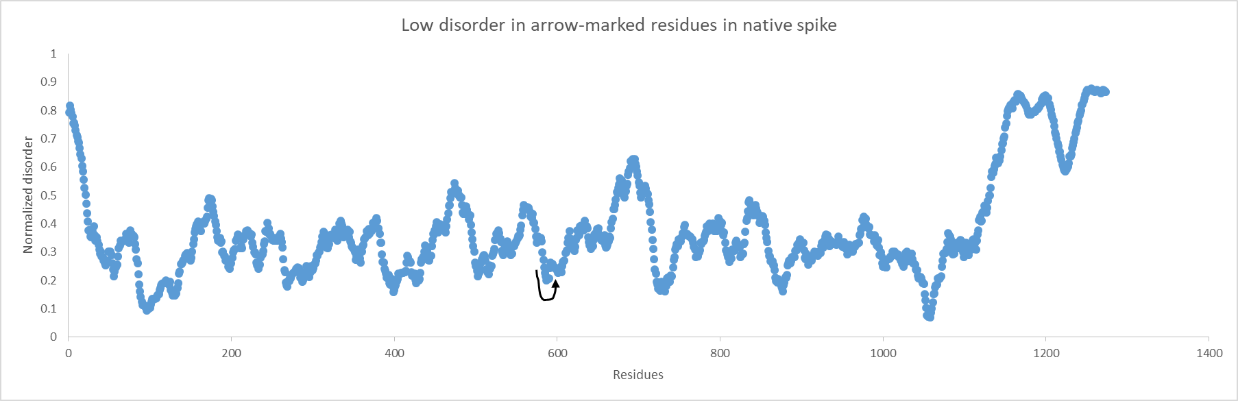
 **C)**

**Fig S11: Disorder plots calculated from AlphaFold models indicate increased fluctuations in remaining potential ADE causing region after deletion.** Increased disorder shown by arrow in **A)** vaccine construct with transmembrane domain and **B)** vaccine construct without transmembrane domain while **C)** less disorder is seen in intact ADE region in native spike with transmembrane domain. The disorder per residue were predicted by AlphaFold-disorder package and normalized per 25 residues’ frame. The graphs were plotted in MS-Office Excel (2013).
